# The synergistic actions of hydrolytic genes reveal the mechanism of *Trichoderma harzianum* for cellulose degradation

**DOI:** 10.1101/2020.01.14.906529

**Authors:** Déborah Aires Almeida, Maria Augusta Crivelente Horta, Jaire Alves Ferreira Filho, Natália Faraj Murad, Anete Pereira de Souza

## Abstract

Bioprospecting genes and proteins related to plant biomass degradation is an attractive approach for the identification of target genes for biotechnological purposes, especially genes with potential applications in the biorefinery industry that can enhance second-generation ethanol production technology. *Trichoderma harzianum* is a potential candidate for cellulolytic enzyme prospection and production. Herein, the enzymatic activities, transcriptome, exoproteome, and coexpression networks of the *T. harzianum* strain CBMAI-0179 were examined under biomass degradation conditions. We used RNA-Seq to identify differentially expressed genes (DEGs) and carbohydrate-active enzyme (CAZyme) genes related to plant biomass degradation and compared them with the genes of strains from congeneric species (*T. harzianum* IOC-3844 and *T. atroviride* CBMAI-0020). *T. harzianum* CBMAI-0179 harbors strain- and treatment-specific CAZyme genes and transcription factors. We detected important proteins related to biomass degradation, including β-glucosidases, endoglucanases, cellobiohydrolases, lytic polysaccharide monooxygenases, endo-1,4-β-xylanases and β-mannanases, in the exoproteome under cellulose growth conditions. Coexpression networks were constructed to explore the relationships among the genes and corresponding secreted proteins that act synergistically for cellulose degradation. An enriched cluster with degradative enzymes was described, and the subnetwork of CAZymes revealed strong correlations among the secreted proteins (AA9, GH6, GH10, GH11 and CBM1) and differentially expressed CAZyme genes (GH45, GH7, AA7 and GH1). Our results provide valuable information for future studies on the genetic regulation of plant cell wall-degrading enzymes. This knowledge can be exploited for the improvement of enzymatic reactions in biomass degradation for bioethanol production.

**Key points:** Different biotechnological approaches were used to understand the mechanism of cellulose degradation of *Trichoderma* spp.

*T. harzianum* CBMAI-0179 is a potential candidate for the production of cellulolytic enzymes.

Coexpression networks revealed genes and proteins acting synergistically for cellulose hydrolysis.

## Introduction

The expanding worldwide demand for renewable and sustainable energy sources has increased the interest in alternative energy sources, and the production of second-generation biofuels seems to be a viable option to confront these issues (dos Santos Castro et al. 2014). Lignocellulosic biomass is the most abundant renewable organic carbon resource on earth, consisting of three major polymers, cellulose, hemicellulose, and lignin (Bayer et al. 2007; dos Santos Castro et al. 2014). However, due to its recalcitrant characteristics, degrading this complex matrix requires a variety of enzymes acting in synergy. Interactions between different enzymes have been investigated to identify optimal combinations and ratios of enzymes for efficient biomass degradation, which are highly dependent on the properties of the lignocellulosic substrates and the surface structure of cellulose microfibrils (van Dyk and Pletschke 2012; Wang et al. 2016). Microorganisms are considered natural producers of enzymes, and many members of both bacteria and fungi have evolved to digest lignocellulose (Ahmed et al. 2017).

Filamentous fungi, including the genera *Trichoderma, Aspergillus, Penicillium* and *Neurospora*, produce extracellular proteins capable of degrading plant cell walls and are widely used in the enzymatic industry (Miao et al. 2015). Species in the filamentous ascomycete genus *Trichoderma* are among the most commonly isolated saprotrophic fungi and are important from a biotechnological perspective (Horta et al. 2018). Most species from this genus are used in agriculture as biocontrol agents due to their ability to antagonize plant-pathogenic fungi, described as mycoparasites, and in industry as producers of hydrolytic enzymes (Druzhinina et al. 2011; Kubicek et al. 2011). Within the *Trichoderma* genus, *T. reesei* is the most intensively studied species and a well-known producer of cellulase and hemicellulase, which are widely employed in industry (Druzhinina and Kubicek 2017; Martinez et al. 2008; Peterson and Nevalainen 2012). However, studies on *T. harzianum* strains have shown their potential to produce different sets of enzymes that can degrade lignocellulosic biomass (Benoliel et al. 2013; Delabona et al. 2012, 2013; Horta et al. 2014); therefore, *T. harzianum* strains are being investigated as potentially valuable sources of new industrial cellulases (Rocha et al. 2016).

The high degree of genetic variation observed within *Trichoderma* species and strains (Al-Sadi et al. 2015; Kubicek et al. 2019) makes it necessary to explore the genetic background of potential *Trichoderma* strains. In the present study, we compared the transcriptome and exoproteome of the potential hydrolytic strain *T. harzianum* CBMAI-0179 with those of *T. harzianum* IOC-3844 and *T. atroviride* CBMAI-0020. Coexpression networks provided novel information about the synergistic action of the hydrolytic enzymes of *T. harzianum* CBMAI-0179. Our findings provide insights into the genes/proteins that act synergistically in plant biomass conversion and can be exploited to improve enzymatic hydrolysis, thereby increasing the efficiency of the saccharification of lignocellulosic substrates for bioethanol production.

## Materials and methods

### Fungal strains, fermentation and enzymatic activities

The species originated from the Brazilian Collection of Environment and Industry Microorganisms (CBMAI, Campinas, SP, Brazil) and the Institute Oswaldo Cruz (IOC, Rio de Janeiro, RJ, Brazil). *T. harzianum* CBMAI-0179 (Th0179), *T. harzianum* IOC-3844 (Th3844) and *T. atroviride* CBMAI-0020 (Ta0020) strains were cultivated on solid medium for 8 days at 37 °C to produce a sufficient number of spores as an inoculum for fermentation, as described in a previous study (Horta et al. 2018). The fermentation process was initiated with the inoculation of 10^7^ spores/mL in an initial volume of 200 mL of Mandels Andreotti (MA) minimal medium (Mandels and Andreotti 1978), and it was performed in biological triplicates using crystalline cellulose (Celuflok, São Paulo, Brazil; degree of crystallinity, 0.72 g/g; composition, 0.857 g/g cellulose and 0.146 g/g hemicellulose) or glucose (control condition) as the unique carbon source. The fermentation process continued for 96 h, and then, the supernatant and RNA samples were collected (Horta et al. 2018).

Supernatants were collected to measure the enzymatic activity (Fig. S1) and determine the exoproteome profile. Xylanase and β-glucosidase activities were determined using the methods described by Bailey and Poutanen (1989) and Zhang et al. (2009), respectively. Cellulase activity was determined using the filter paper activity (FPA) test according to Ghose (1987). Protein levels were measured based on the Bradford (1976) method.

### RNA extraction

Mycelial samples from both conditions were extracted from Th0179, Th3844 and Ta0020, stored at −80 °C, ground in liquid nitrogen using a mortar and pestle, and used for RNA extraction using the LiCl RNA extraction protocol (Oliveira et al. 2015).

### Library construction and RNA-Seq

RNA samples were quantified using a NanoDrop 8000 (Thermo Scientific, Wilmington, DE, USA). Paired-end libraries were constructed using 1 µg of each RNA sample obtained from the mycelial samples and the TruSeq RNA Sample Preparation Kit v2 (Illumina 2014) (Illumina Inc., San Diego, CA, USA) according to the manufacturer’s instructions. The expected target sizes were confirmed using a 2100 Bioanalyzer (Agilent Technologies, Palo Alto, CA, USA) and the DNA 1000 Kit, and the libraries were quantified by qPCR using the KAPA library quantification Kit for Illumina platforms (Kapa Biosystems, Wilmington, MA, USA). The average insertion size was 260 bp. The sequencing was carried out on the HiSeq 2500 platform (Illumina, San Diego, CA, USA) according to the manufacturer’s specifications for paired-end reads of 150 bp.

### Data sources

The reads were deposited into the Sequence Read Archive (SRA) database in the National Center for Biotechnology Information (NCBI) under BioProject number PRJNA336221. The nucleotide and protein sequences of *T. harzianum* T6776 (PRJNA252551) (Baroncelli et al. 2015) and *T. atroviride* IMI 206040 (PRJNA19867) (Kubicek et al. 2011) were used as a reference for transcriptome mapping and downloaded from the NCBI database (www.ncbi.nlm.nig.gov).

### *Phylogenetic analysis of* Trichoderma *spp*

Amplification of the internal transcribed spacer (ITS) region from the genomic DNA of the strains under study was conducted via PCR. Multiple sequence alignment was performed using ClustalW (Thompson et al. 1994), and a phylogenetic tree was created using the Molecular Evolutionary Genetics Analysis v7.0 (MEGA7) program (Kumar et al. 2016). The maximum likelihood (ML) (Jones et al. 1992) method of inference was used based on the Kimura two-parameter (K2P) model (Kimura 1980). We used 1,000 bootstrap replicates (Felsenstein 1985) for each analysis. The trees were visualized and edited using the FigTree program v1.4.3 (http:/tree.bio.ed.ac.uk/software/figtree/). The phylogenetic tree showed a high phylogenetic proximity between Th0179 and Th3844 (with bootstrap support of 92%), while Ta0020 shared closer affinity with *T. atroviride* CBS 142.95 (Fig. S2) (Rosolen et al. 2020).

### Transcriptome analysis

The quality of the sequencing reads was assessed with FastQC v0.11.5 (Andrews 2010), and the trimming with a sliding window of size 4, minimum quality of 15, and length filtering (minimal length of 36 bp) was performed with Trimmomatic v0.36 (Bolger et al. 2014). The RNA-Seq data were analyzed using CLC Genomics Workbench software (v6.5.2; CLC bio, Finlandsgade, Denmark) (CLC Genomics Workbench 2016). The reads were mapped against the reference genomes of *T. harzianum* T6776 (Baroncelli et al. 2015) and *T. atroviride* IMI 206040 (Kubicek et al. 2011) using the following parameters: minimum length fraction = 0.5; minimum similarity fraction = 0.8; and maximum number of hits for a read = 10. For the paired settings, the parameters were minimum distance = 150 and maximum distance = 300, including the broken pairs. The gene expression values were normalized in transcripts per million (TPM) (Conesa et al. 2016). To statistically analyze the differentially expressed genes (DEGs), fold change ≥ 1.5 or ≤ -1.5 and p-value ≤ 0.05 were applied. Venn diagrams were constructed to compare the genes with TPM expression values greater than zero under both conditions from all strains (http://bioinformatics.psb.ugent.be/webtools/Venn/).

### Transcriptome annotation and CAZyme determination

Sequences were functionally annotated according to the Gene Ontology (GO) terms (Ashburner et al. 2000) with Blast2Go v4.1.9 (Conesa et al. 2005) using BLASTx-fast and a cutoff e-value of 10^−6^. Information derived from the CAZy database (Lombard et al. 2014) was downloaded (www.cazy.org) to locally build a CAZy database. The protein sequences of *T. harzianum* T6776 and *T. atroviride* IMI 206040 were used as queries in basic local alignment search tool (BLAST) searches against the locally built CAZy database. BLAST matches showing an e-value less than 10^−11^, identity greater than 30% and queries covering greater than 70% of the sequence length were selected and classified according to the CAZyme catalytic group as follows: glycoside hydrolases (GHs), carbohydrate-binding modules (CBMs), glycosyltransferases (GTs), carbohydrate esterases (CEs), auxiliary activities (AAs) or polysaccharide lyases (PLs). CAZymes were also annotated according to their Enzyme Commission (EC) number (Yamanishi et al. 2009) through the Braunschweig Enzyme Database (BRENDA) (Schomburg et al. 2017) (www.brenda-enzymes.org) using BLASTp with an e-value cutoff of 10^−10^, identity greater than 30% and queries covering greater than 60% of the sequence length.

### Coexpression networks

For the coexpression networks of Th0179, TPM expression values were used to perform the analyses. The network was assembled by calculating Pearson’s correlation coefficient for each pair of genes. Genes showing null values for most of the replicates under different experimental conditions were excluded. The cluster analysis procedure was performed with the heuristic cluster chiseling algorithm (HCCA), and the highest reciprocal rank (HRR) (Mutwil et al. 2011; Mutwil et al. 2010) was used to empirically filter the edges, retaining edges with a HRR ≤ 30. Cytoscape software v3.6.0 (Shannon et al. 2003) was used for network visualization.

### Exoproteome analysis

The analysis of the exoproteome of Th0179 under both fermentative conditions was performed via liquid chromatography tandem mass spectrometry (LC-MS/MS) using the data-independent method of acquisition MS^E^ as described by Horta et al. (2018). The LC-MS/MS data were processed using ProteinLynx Global Server (PLGS) v3.0.1 software (Waters, Milford, MA, USA), and the proteins were identified by comparison to the *Trichoderma* sequence database available in the UniProt Knowledgebase (UniProtKB; https://www.uniprot.org/uniprot/). BLASTp searches of the fasta sequences of the identified proteins were performed against the *T. harzianum* T6776 genome to identify the secreted proteins and compare them to the transcriptome.

### RT-qPCR analysis

To verify the reliability and accuracy of the transcriptome data, reverse transcription-quantitative PCR (RT-qPCR) was performed to validate the differential expression results. The RNeasy Mini Kit (Qiagen, Hilden, Germany) was used for RNA extraction, and cDNA was synthesized using the QuantiTect Reverse Transcription Kit (Qiagen, Hilden, Germany). Primers were designed using the Primer3Plus (Untergasser et al. 2007); the target amplicon sizes ranged from 120-200 bp, with an optimal annealing temperature of 60 °C and an optimal primer length of 20 bp.

Quantification of gene expression was performed by continuously monitoring SYBR Green fluorescence. The reactions were performed in triplicate in a total volume of 6.22 μL. Each reaction contained 3.12 μL of SYBR Green Supermix (Bio-Rad, Hercules, CA, USA), 1.0 μL of forward and reverse primers and 2.1 μL of diluted cDNA. The reactions were assembled in 384-well plates. PCR amplification-based expression profiling of the selected genes was performed using specific endogenous controls for each strain (Table S1). RT-qPCR was conducted with the CFX384 Touch Real-Time PCR Detection System (Bio-Rad). Gene expression was calculated via the delta-delta cycle threshold method (Livak and Schmittgen 2001). The obtained RT-qPCR results were compared with the RNA-Seq results. The fold changes of the selected genes exhibited the same expression profiles between the RT-qPCR and RNA-Seq analyses (Fig. S3).

## Results

### Transcriptome analysis of *Trichoderma spp*. under cellulose growth conditions

The present study represents the first deep genetic analysis of Th0179, describing and comparing the transcriptome under two different growth conditions, cellulose and glucose, to identify the genes involved in plant biomass degradation. Reads were mapped against reference genomes of *T. harzianum* (PRJNA252551) and *T. atroviride* (PRJNA19867), generating 96.3, 111.8 and 133.3 million paired-end reads for Th0179, Th3844 and Ta0020, respectively. To establish the degrees of similarity and difference in gene expression among strains and between treatments, a principal component analysis (PCA) was performed using the *T. harzianum* T6776 genome as a reference. The PCA results showed clustered groups with higher similarity between treatments than among strains. The transcriptomes of Th0179 and Th3844 were more similar to each other than to the transcriptome of Ta0020 (Fig. 1a), showing that it is possible to capture differences among the three strains.

**Fig. 1.**
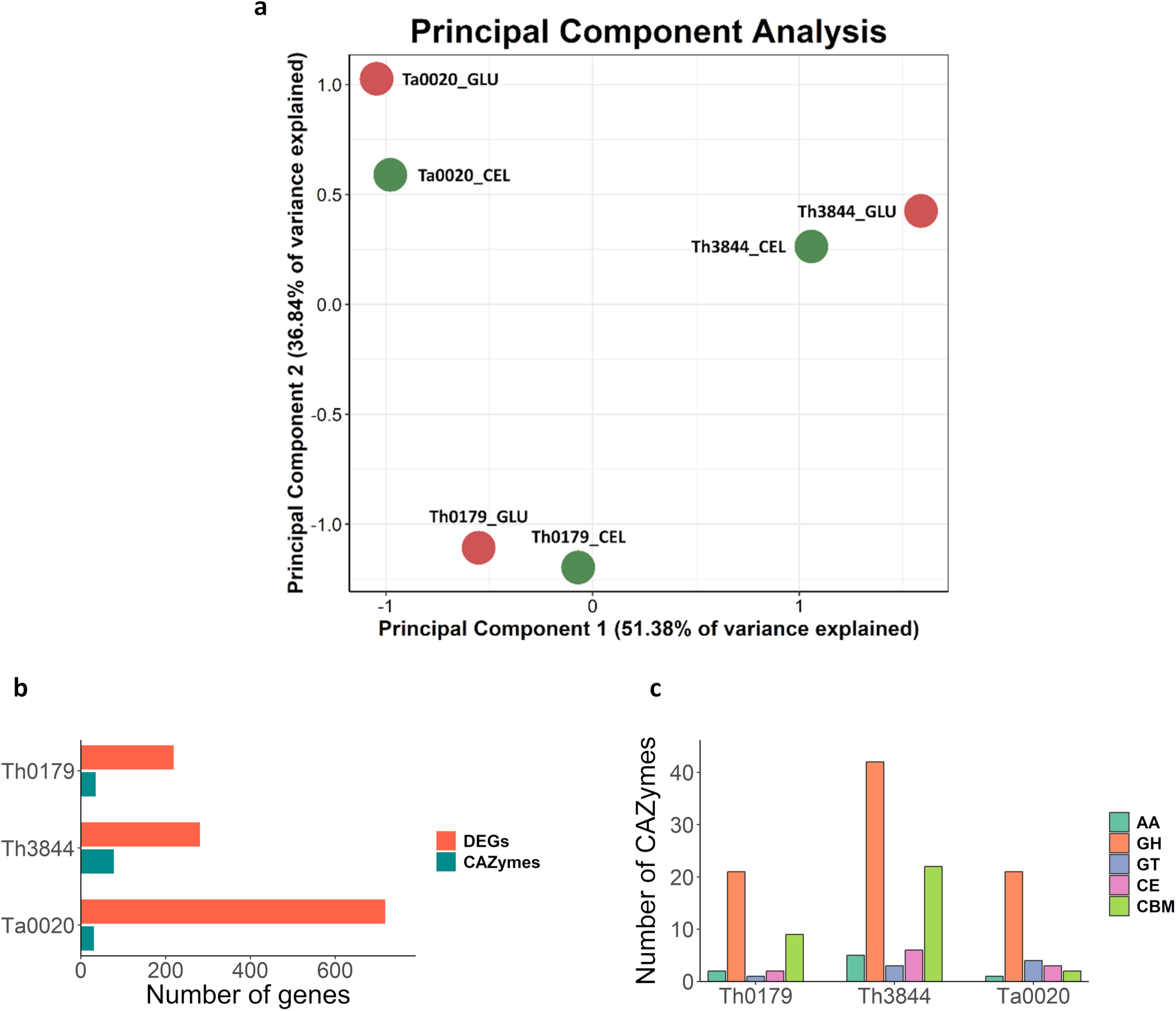
The transcriptome profiles, gene expression comparison and the main CAZyme classes identified for all strains. PCA of transcriptome mapping according to species and growth conditions (CEL – cellulose and GLU – glucose) using the *T. harzianum* T6776 genome as a reference (a), number of DEGs and differentially expressed CAZyme genes in *T. harzianum* CBMAI-0179 (Th0179), *T. harzianum* IOC-3844 (Th3844), and *T. atroviride* CBMAI-0020 (Ta0020) under cellulose growth conditions (b), the differentially expressed CAZyme classes identified for each strain under cellulose growth conditions (c)

Venn diagrams of the genes exhibiting expression levels greater than zero were constructed based on the similarities among the genes from all strains, showing strain-specific genes expressed under both conditions (Fig. S4). The number of genes shared by Th0179 and Th3844 was higher under both conditions than that shared by either Th0179 or Th3844 and Ta0020. Th0179 exhibited the highest number of unique expressed genes, with 374 and 168 unique genes under the cellulose and glucose growth conditions, respectively, in which major facilitator superfamily (MFS) and ATP binding cassette (ABC) transporters and transcription factors were found (Table S2). Among the genes that were upregulated in cellulose conditions relative to glucose conditions, 219 were identified for Th0179, 281 were identified for Th3844, and 718 were identified for Ta0020 (Fig. 1b and Table S3).

### CAZyme identification and distribution in *Trichoderma spp*

The set of CAZymes was identified by mapping all of the protein sequences from *T. harzianum* T6776 and *T. atroviride* IMI 206040 against the CAZy database. Based on the filtering criteria, a total of 631 proteins were retained as CAZyme genes for *T. harzianum*, which corresponds to 5.5% of the total 11,498 proteins predicted for this organism (Baroncelli et al. 2015), and 640 proteins were retained for *T. atroviride*, which corresponds to 5.4% of the total 11,816 proteins predicted for this organism (Kubicek et al. 2011).

Considering the DEGs under cellulose growth conditions, we found 35, 78 and 31 differentially expressed CAZyme genes for Th0179, Th3844 and Ta0020, respectively (Fig. 1b). The contents of the main CAZyme classes (AA, GH, GT, CE and CBM) were identified (Fig. 1c), and the differences regarding the specific CAZyme families for each strain are shown in Fig. 2. The classification of the CAZyme genes along with their enzyme activities, fold change values, e-values, EC numbers and TPM expression values for Th0179, Th3844 and Ta0020 under cellulose fermentative conditions are described in Table S4.

**Fig. 2.**
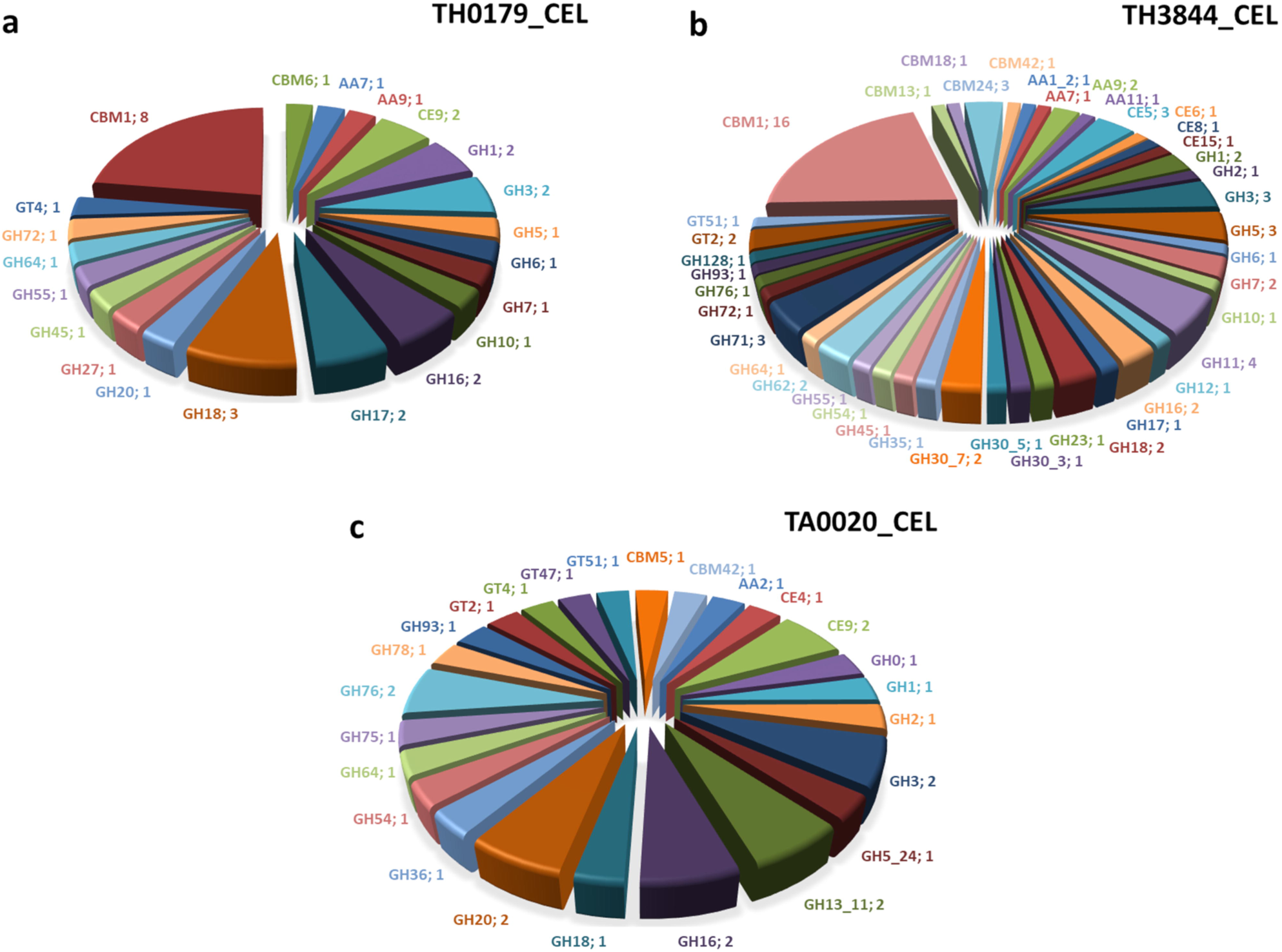
Distribution of CAZyme families in *Trichoderma* spp. Classification and quantification of CAZyme families in *T. harzianum* CBMAI-0179 (a), *T. harzianum* IOC-3844 (b), and *T. atroviride* CBMAI-0020 (c) under cellulose growth conditions

The expression levels of the main cellulase and hemicellulase families based on their principal enzyme activity present in Th0179, Th3844 and Ta0020 were evaluated using the transcriptomic data (Fig. 3). GHs related to cellulose degradation were found in Th3844, including two AA9 members with a CBM1 module for cellulose binding. All GHs except GH12, including AA9/CBM1, were detected in Th0179. Only one gene belonging to the GH5 family (TRIATDRAFT_81867 – 48.12 TPM) with cellulase activity was observed in Ta0020. The most highly expressed genes were cellobiohydrolases from the GH6/CBM1 (THAR02_04414 – 299.66 TPM for Th0179 and 986.25 TPM for Th3844) and GH7/CBM1 families (THAR02_08897 – 369.92 TPM for Th0179; THAR02_03357 and THAR02_08897 – 1876.89 TPM for Th3844) (Fig. 3a).

**Fig. 3.**
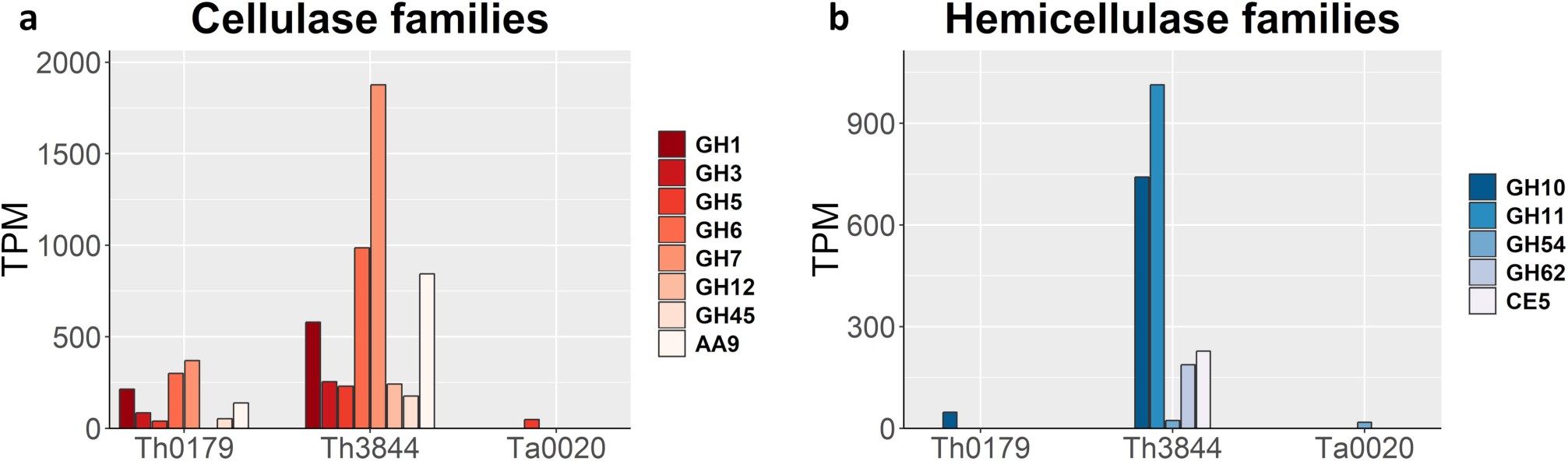
Evaluation of CAZyme family expression in *Trichoderma* spp. via RNA-Seq. Quantification of the expression of the main families related to cellulose (a) and hemicellulose (b) degradation in TPM

In addition, families related to hemicellulose degradation were identified in Th3844, in which endo-1,4-β-xylanases from the GH11 family had the greatest expression levels (THAR02_02147, THAR02_05896, THAR02_08630 and THAR02_08858 – 1013.62 TPM). Endo-1,4-β-xylanase from the GH10 family was found in both *T. harzianum* strains (THAR02_03271 – 47.58 TPM for Th0179 and 741.35 TPM for Th3844). Acetylxylan esterases from the CE5/CBM1 family were only found in Th3844 (THAR02_01449 and THAR02_07663 – 227.56 TPM). We identified only one α-L-arabinofuranosidase from the GH54/CBM42 family (TRIATDRAFT_81098 – 18.07 TPM) in Ta0020, whereas for Th3844, three α-L-arabinofuranosidases from the GH54/CBM42 and GH62 families were identified (Fig. 3b). Despite the higher number of DEGs, the Ta0020 strain showed the lowest number and expression levels of CAZyme genes related to cellulose and hemicellulose degradation.

### Functional annotation of *T. harzianum* CBMAI-0179 in the presence of cellulose

The first functional annotation of the expressed genes under cellulose fermentative conditions for Th0179 was performed (Fig. 4). A total of 7,718 genes were annotated, which corresponds to 67.1% of the total genes predicted for the reference strain *T. harzianum* T6776 (Baroncelli et al. 2015). Compared with Th3844, Th0179 presented 163 more genes related to catalytic activity, 53 more genes related to hydrolase activity, and 15 more genes associated with TF activity but 10 fewer genes related to transmembrane transport, suggesting that these two strains have developed different functional regulation. Ta0020 presented fewer genes related to any of the above functions than Th3844 and Th0179 except for TF activity, for which Ta0020 had 53 more genes than Th0179.

**Fig. 4.**
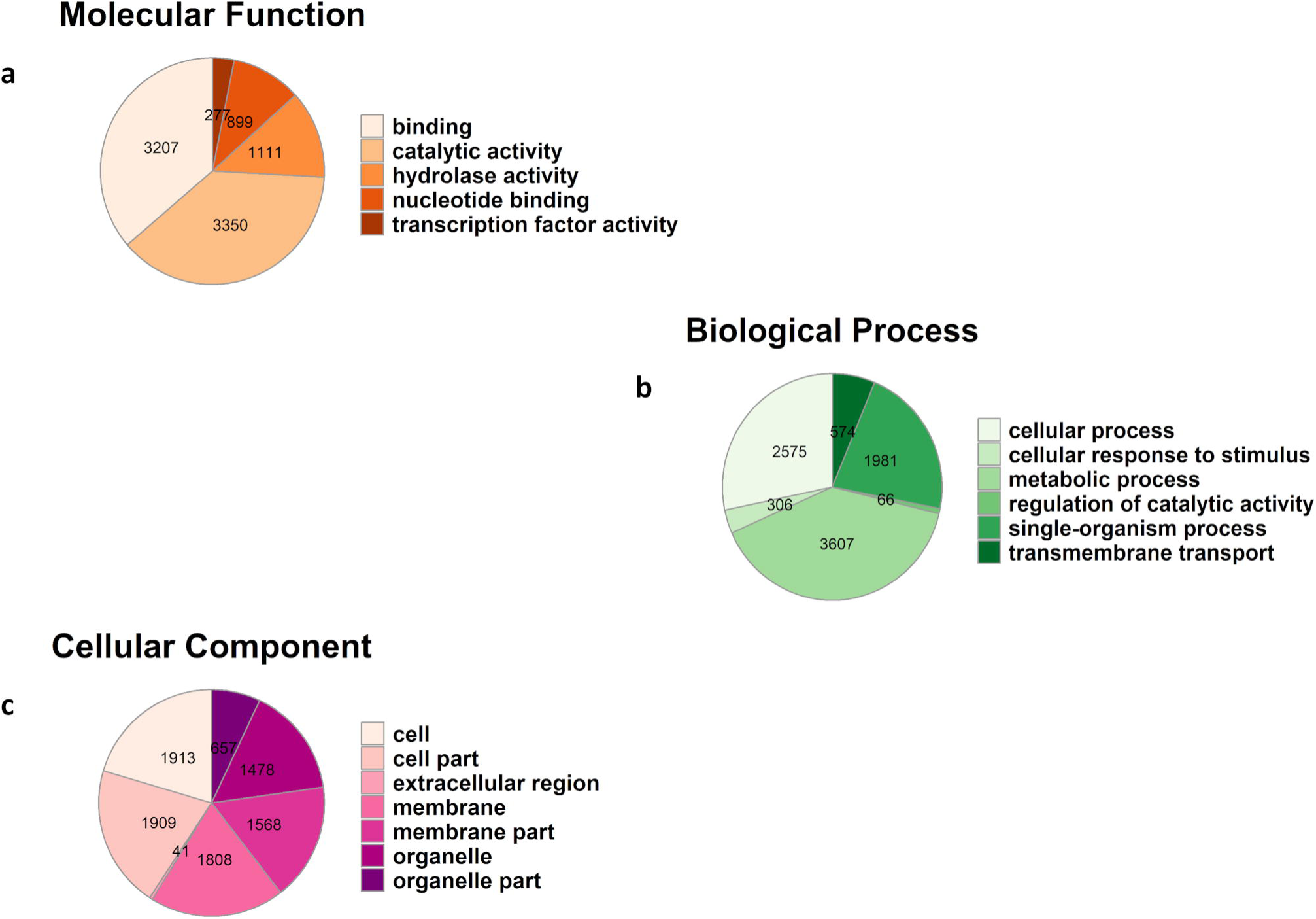
GO terms of *T. harzianum* CBMAI-0179 under cellulose growth conditions. The genes were annotated according to the main GO terms: molecular function (a), biological process (b), and cellular component (c)

### Exoproteome and RNA-Seq data correlation of *T. harzianum* CBMAI-0179

Once the transcriptome was characterized, we analyzed the secreted proteins identified in the exoproteome profile of Th0179. We identified 32 proteins in the cellulose aqueous extract, 12 only in the glucose aqueous extract, and 20 under both conditions (Table S5). Among the 32 secreted proteins detected using cellulose as the carbon source, the main CAZyme families observed were β-glucosidases (EC 3.2.1.21), endo-β-1,4-glucanases (EC 3.2.1.4), lytic polysaccharide monooxygenases (LPMOs) (EC 3.2.1.4), cellobiohydrolases (EC 3.2.1.91), endo-1,4-β-xylanases (EC 3.2.1.8), and β-mannanases (EC 3.2.1.78). Two genes from the GH3 (THAR02_00656 and THAR02_00890) and GH5/CBM1 (THAR02_04405 and THAR02_09719) families and one gene from the AA9/CBM1 (THAR02_02134) and GH6/CBM1 (THAR02_04414) families corresponded to the main group of secreted cellulases, whereas one gene from the GH10 (THAR02_03271) family and two genes from the GH11 (THAR02_02147 and THAR02_05896) family corresponded to the main group of hemicellulases detected in the supernatant. In addition, a member of the GH5 cellulase family with β-mannanase activity (THAR02_03851) was identified, and 8 other proteins were classified as uncharacterized proteins. We also identified hemicellulases in both extracts, such as α-L-arabinofuranosidase B (EC 3.2.1.55) and xylan 1,4-β-xylosidase (EC 3.2.1.37).

Correlating the exoproteome with the transcriptome data under cellulose growth conditions, we observed CAZyme genes showing high TPM values in the cellulose condition, including cellulose 1,4-β-cellobiosidase (EC 3.2.1.91) from the GH6/CBM1 family (299.66 TPM), cellulase (EC 3.2.1.4) from the AA9/CBM1 family (138.62 TPM) and endo-1,4-β-xylanase (EC 3.2.1.8) from the GH11/CBM1 family (119.56 TPM). Two uncharacterized proteins (THAR02_02133 and THAR02_08479) with the CBM1 module of cellulose binding also showed increased expression levels under the cellulose condition. In contrast, the THAR02_00656 gene, which displays β-glucosidase (EC 3.2.1.21) activity, from the GH3 family had the lowest expression level (5.08 TPM) among the CAZyme genes, indicating that genes with low expression levels are also important for functional secreted proteins (Alfaro et al. 2016).

### Coexpression networks for *T. harzianum* CBMAI-0179

The coexpression network was assembled for Th0179 and included all genes obtained from the mapping results using the *T. harzianum* T6776 genome as a reference under both conditions. The DEGs and CAZyme genes were highlighted in the network (Fig. 5a), composed of a total of 11,104 nodes with 153,893 edges. We detected 219 genes corresponding to the DEGs under cellulose growth conditions and 367 DEGs under glucose growth conditions. Differentially expressed CAZyme genes from both conditions were also identified in this network, with 35 and 28 CAZymes under cellulose and glucose growth conditions, respectively. The DEGs in glucose clustered on the top while DEGs in cellulose clustered on the bottom part of the network, reflecting the different regulation of degradative activities according to the substrate.

**Fig. 5.**
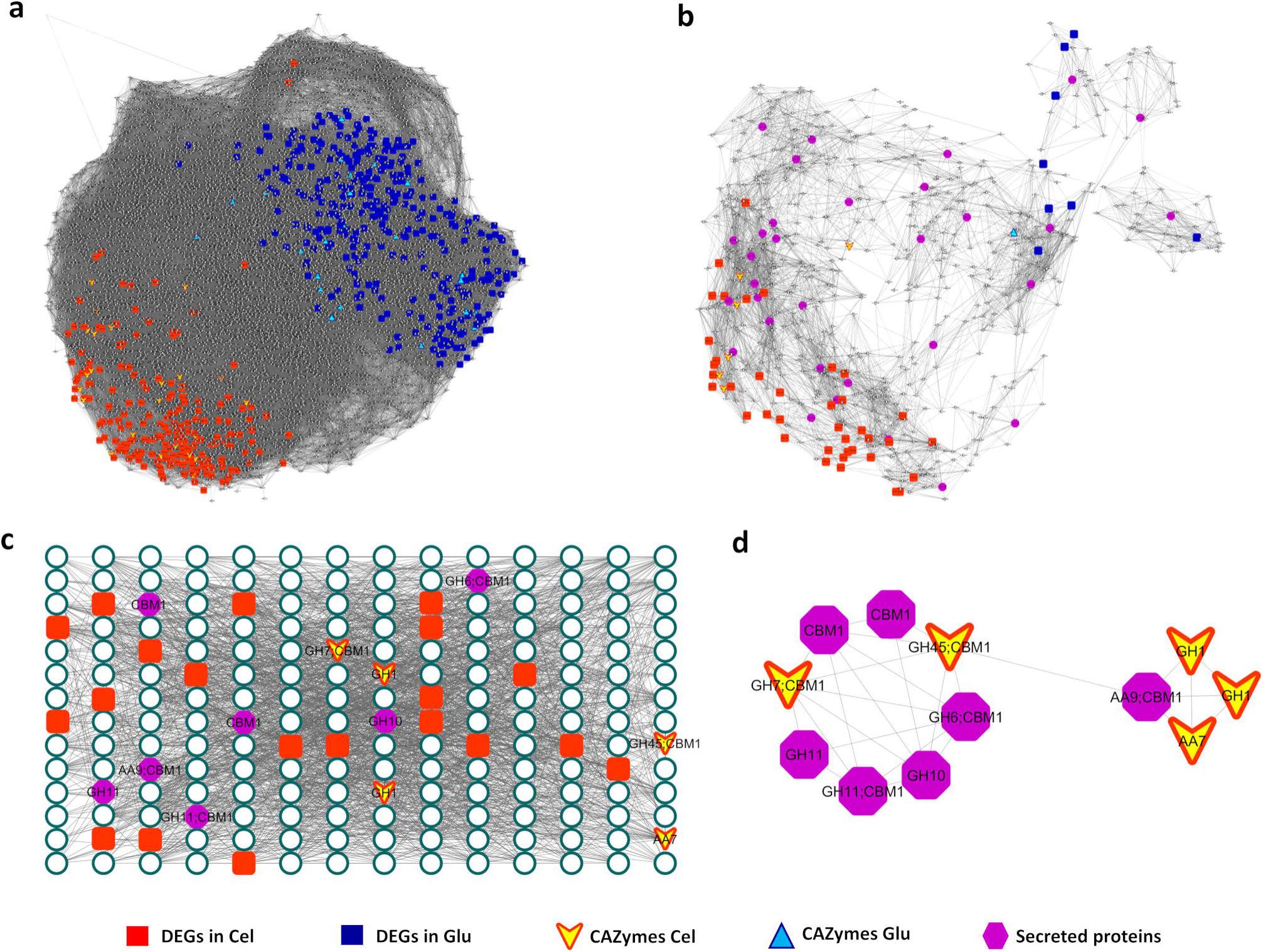
Coexpression networks of *T. harzianum* CBMAI-0179. Complete coexpression network (a), the coexpression subnetwork based on the exoproteome data (b), the enriched cluster analysis of the coexpression network (c), the subnetwork of the CAZyme genes and the secreted proteins identified in the cluster analysis (d). Red squares indicate DEGs under cellulose growth conditions, blue squares indicate DEGs under glucose growth conditions, yellow triangles indicate CAZyme genes under cellulose growth conditions, light blue triangles indicate CAZyme genes under glucose growth conditions and purple hexagons indicate the secreted proteins under cellulose growth conditions

A subnetwork was generated based only on the secreted proteins that were present in the cellulose aqueous extract and that had corresponding genes in the coexpression network (Fig. 5b). This subnetwork exclusively represented the genes and their closest related genes that are correlated with the secreted proteins. It was composed of 713 nodes and 6,124 edges, and includes the 32 genes that encode the secreted proteins. The CAZyme genes found in this subnetwork included the GH1 family (THAR02_02251 and THAR02_05432) with β-glucosidase activity, the GH7/CBM1 family (THAR02_08897) with cellulose 1,4-β-cellobiosidase activity, and the GH45/CBM1 family (THAR02_02979) with cellulase activity. Despite the different functions of the correlated genes, it is predicted that these genes participate in the genetic regulation of the detected CAZymes/proteins and are important to the regulation of the hydrolytic system.

The cluster analysis classified 11,102 genes in 84 clusters (Table S6). We identified an enriched cluster composed of 196 nodes and 2,125 edges (Fig. 5c and Table S6) containing the greatest number of CAZyme genes among the clusters. Of the 12 CAZyme genes detected, 7 corresponded to the secreted proteins detected in the cellulose aqueous extract, and 5 were differentially expressed. In addition, we identified 20 DEGs under cellulose growth conditions and 119 uncharacterized proteins in this cluster, showing a strong correlation with the CAZyme genes, which indicates unknown genes that are important to the degradation process. The CAZyme genes from the cluster analysis were selected to generate a new subnetwork with their corresponding edges (Fig. 5d), showing linear correlations among secreted proteins (AA9, GH6, GH10, GH11 and CBM1) and differentially expressed CAZyme genes (GH45, GH7, AA7 and GH1).

## Discussion

In this study, different biotechnological approaches were used in bioprospecting new and efficient enzymes for possible applications in the enzymatic hydrolysis process. We performed the enzymatic activity, transcriptome, exoproteome and coexpression network analyses of the Th0179 strain, which has potential for plant biomass degradation under cellulose growth conditions, to gain insight into the genes and proteins produced that are associated with cellulose hydrolysis. Furthermore, we compared the expression levels with *Trichoderma* spp. that have been previously studied under the same conditions (Horta et al. 2018) to understand how the regulation varies between hydrolytic strains (*T. harzianum* CBMAI-0179 and *T. harzianum* IOC-3844) and a common mycoparasitic ancestor species (*T. atroviride* CBMAI-0020).

Through the transcriptome analysis, we identified the DEGs and the differentially expressed CAZyme genes in all strains. Although the Ta0020 strain presented the highest number of identified DEGs, it was found to be less efficient than other *Trichoderma* strains in degrading plant biomass. *T. atroviride* is mostly used as a biocontrol agent and is among the best mycoparasitic fungus used in agriculture (Kubicek et al. 2011). Each strain presented a different set of genes with different expression levels, which can be attributed to strain differences in the regulatory mechanisms of hydrolysis. Additionally, differences regarding the carbon source metabolization were observed, which plays an important role in the production of enzymes and promotes the expression of different sets of genes as the fungus seeks to adapt to new environments (Horta et al. 2018). Even though all species are induced by cellulose growth, *T. harzianum* strains showed high numbers of CAZyme genes with enhanced specificity for that type of biomass.

Among the main CAZyme classes detected in all strains, GHs were well represented, with 21 genes in Th0179 and twice that in Th3844. GHs compose an extremely important class in several metabolic routes in fungi, including genes involved in cellulose and chitin degradation (Ferreira Filho et al. 2017). Here, we found the main CAZyme families responsible for cellulose degradation, such as GH1, GH3, GH5, GH6, GH7, GH12, GH45 and an LPMO from the AA9 family, and for hemicellulose degradation, such as GH10, GH11, GH54, GH62 and CE5 (Ferreira Filho et al. 2017; Sweeney and Xu 2012; Villares et al. 2017; Zhao et al. 2014). Within the GH class, an important family expected in our data was GH18, which has chitinase activity; since members of the genus *Trichoderma* (*T. harzianum* and *T. atroviride*) are capable of mycoparasitism, this class is directly related to the biological control of these species (Binod et al. 2007; Martinez et al. 2008).

In comparing the two *T. harzianum* strains (Th0179 and Th3844), we observed that Th0179 presented fewer CAZyme genes related to cellulose degradation than Th3844, with lower expression levels (Fig. 3a). However, the measurement of the enzymatic activity from the culture supernatants after 96 h of growth showed similar cellulase activity profiles during growth on cellulose, suggesting a greater potential of Th0179 to degrade cellulose through the use of a more efficient set of enzymes. A similar profile was observed in *T. reesei*, the most studied fungus within this genus and an important industrial producer of cellulolytic enzymes (Druzhinina et al. 2011); few CAZyme genes have been detected in its machinery, but it can reach the highest cellulolytic activity (Druzhinina and Kubicek 2017; Horta et al. 2018; Manika et al. 2014). It is interesting to observe which proteins found in the exoproteome of Th0179 may respond to this increased cellulase activity.

CAZyme genes related to cellulose degradation, including endo-β-1,4-glucanases, β-glucosidases, and cellobiohydrolases, and CAZyme genes related to hemicellulose degradation, such as endo-1,4-β-xylanases, were identified in the Th0179 transcriptome. In addition, we verified important secreted CAZymes that play important roles in biomass degradation, such as β-glucosidases (A0A0F9XRC5 and A0A0F9XQT4), LPMO (A0A0F9XMI8), cellulose 1,4-β-cellobiosidase (nonreducing end) (A0A0G0AEM7), cellulases (A0A0F9XG06 and A0A0F9WYH5), endo-1,4-β-xylanases (A0A0F9Y0Y9, A0A0H3UCP8, and A0A0F9XXA4), and endo-1,4-β-mannosidase (A0A0G0AGG8). One β-glucosidase from the GH3 family (THAR02_00656) was found in the exoproteome, which had the lowest expression level among the CAZymes related to biomass degradation; the same pattern was observed for *T. harzianum* IOC-3844 (Horta et al. 2018).

Horta et al. (2018) analyzed the exoproteomes of Th3844 and Ta0020 under equivalent conditions, selecting a set of 80 proteins for a complete classification and analysis of expression levels based on their transcriptomes. Comparison of the Th0179 and Th3844 exoproteomes indicated that both produced many of the main CAZymes listed above; however, the THAR02_05896 gene, which encodes an endo-1,4-β-xylanase protein, and the THAR02_03851 gene, which encodes a protein with endo-1,4-β-mannosidase activity, are exclusively present in the Th0179 exoproteome. In the present study, when these exoproteomes were compared, we observed that most of the proteins were upregulated in cellulose only for Th0179, including the AA9/CBM1 family (THAR02_02134) that showed a 2.65-fold change (138.62 TPM). Among the strains, Ta0020 exhibited the lowest number of secreted proteins related to cellulose and hemicellulose degradation. Thus, the exoproteome analysis identified key enzymes that are fundamental for cellulose hydrolysis and act synergistically for efficient plant biomass degradation.

The organization of the transcriptomic data into coexpression networks using graph theory allowed the construction of gene interaction networks that were represented by nodes connected to edges (Azevedo et al. 2015; Mutwil et al. 2010). Nodes represent the genes, and the edges represent the connections among these genes. Correlations are determined based on the expression level of the genes pair by pair, indicating that genes spatially closer to one another are more highly correlated than the genes that are farther apart. The coexpression subnetwork revealed complex, specific relationships between CAZyme genes and genes involved in the production and secretion of the detected proteins and is helpful for understanding the functions and regulation of genes. We identified an enriched cluster with strong correlations among genes, CAZyme genes, secreted proteins and unknown genes in the cluster analysis. Networks such as this one can be used as a platform to search for target genes or proteins in future studies to comprehend the synergistic relationships between genes, their regulation and protein production, which is very useful information for understanding the saccharification process.

Furthermore, through the functional annotation analysis of Th0179, we identified TFs and transporters, which should be investigated in further studies to better understand the mechanisms by which these genes are regulated. Most sugar transporters have yet to be characterized but play important roles in taking up mono- or disaccharides into fungal cells after biomass degradation (de Gouvea et al. 2018; Peng et al. 2018). Fungal gene expression is controlled at the transcriptional level (Liu et al. 2017), and its regulation affects the composition of enzyme mixtures; accordingly, it is explored in several species because of its potential applications (Benocci et al. 2017). This further emphasizes the importance of deleting and/or overexpressing TFs that regulate specific genes directly involved in plant biomass degradation (Liu et al. 2017).

In summary, the analyses of the transcriptome, exoproteome and coexpression networks revealed important genes and enzymes that *T. harzianum* CBMAI-0179 uses to hydrolyze cellulose and that most likely act synergistically to depolymerize polysaccharides. The results suggest great potential of this strain to degrade cellulose and can contribute to the optimization of enzymatic cocktails for bioethanol production.

## Conclusions

Bioprospecting new catalytic enzymes and improving technologies for the efficient enzymatic conversion of plant biomass are required for advancing biofuel production. *T. harzianum* CBMAI-0179 is a novel potential candidate producer of plant cell wall polysaccharide-degrading enzymes that can be biotechnologically exploited for plant biomass degradation. The cellulase activity profile indicated high efficiency for the set of genes and proteins produced and consequently the potential of this strain for cellulose degradation. A set of highly expressed CAZymes and proteins that are strain- and treatment-specific was described in both the transcriptome and exoproteome analyses. The coexpression network revealed the synergistic gene action to hydrolyze cellulose, in which the cluster analysis revealed genes with strong correlations necessary for saccharification. Combined, these tools provide a powerful approach for catalyst discovery and the selection of target genes for the heterologous expression of proteins. In future studies, these tools can aid the selection of new species and the optimization of the production of powerful enzymes for use in enzymatic cocktails for second-generation bioethanol production.

## Declarations

### Ethical approval

This article does not contain any studies with human participants or animals performed by any of the authors.

### Consent to participate

Not applicable.

### Consent for publication

Not applicable.

### Availability of data and material

The datasets generated and/or analyzed during the current study are included in this published article and its supplementary material. The reads have been deposited at the NCBI SRA and can be accessed under the BioProject number PRJNA336221.

### Authors’ contributions

DAA and APS conceived and designed the study. DAA, MACH, JAFF and NFM performed the data analysis. MACH designed and performed the fermentation experiments. DAA drafted the manuscript, which was critically revised by MACH, JAFF and APS. All authors read and approved the final manuscript.

## Acknowledgments

We would like to acknowledge the funding from Fundação de Amparo à Pesquisa do Estado de São Paulo (FAPESP), Coordenação de Aperfeiçoamento de Pessoal de Nível Superior (CAPES, Computational Biology Program) and Conselho Nacional de Desenvolvimento Científico e Tecnológico (CNPq). We thank the National Institute of Metrology, Quality and Technology (INMETRO) for performing the proteomics analysis via LC-MS/MS, the Brazilian Biorenewables National Laboratory (LNBR), Campinas – SP, for conducting the fermentation experiments and the Center of Molecular Biology and Genetic Engineering (CBMEG) at the University of Campinas, SP, for use of the center and laboratory space. This manuscript was previously posted to bioRxiv https://www.biorxiv.org/content/10.1101/2020.01.14.906529v2

