## Supplementary Material for "The synergistic actions of hydrolytic genes reveal the mechanism of *Trichoderma harzianum* for cellulose degradation"

Prof Anete Pereira de Souza

**ORCID**

0000-0002-5095-5177, 0000-0003-2159-5296, 0000-0001-9690-7704, 0000-0001-7652-2567, 0000-0003-3831-9829

**Journal: “Applied Microbiology and Biotechnology”**

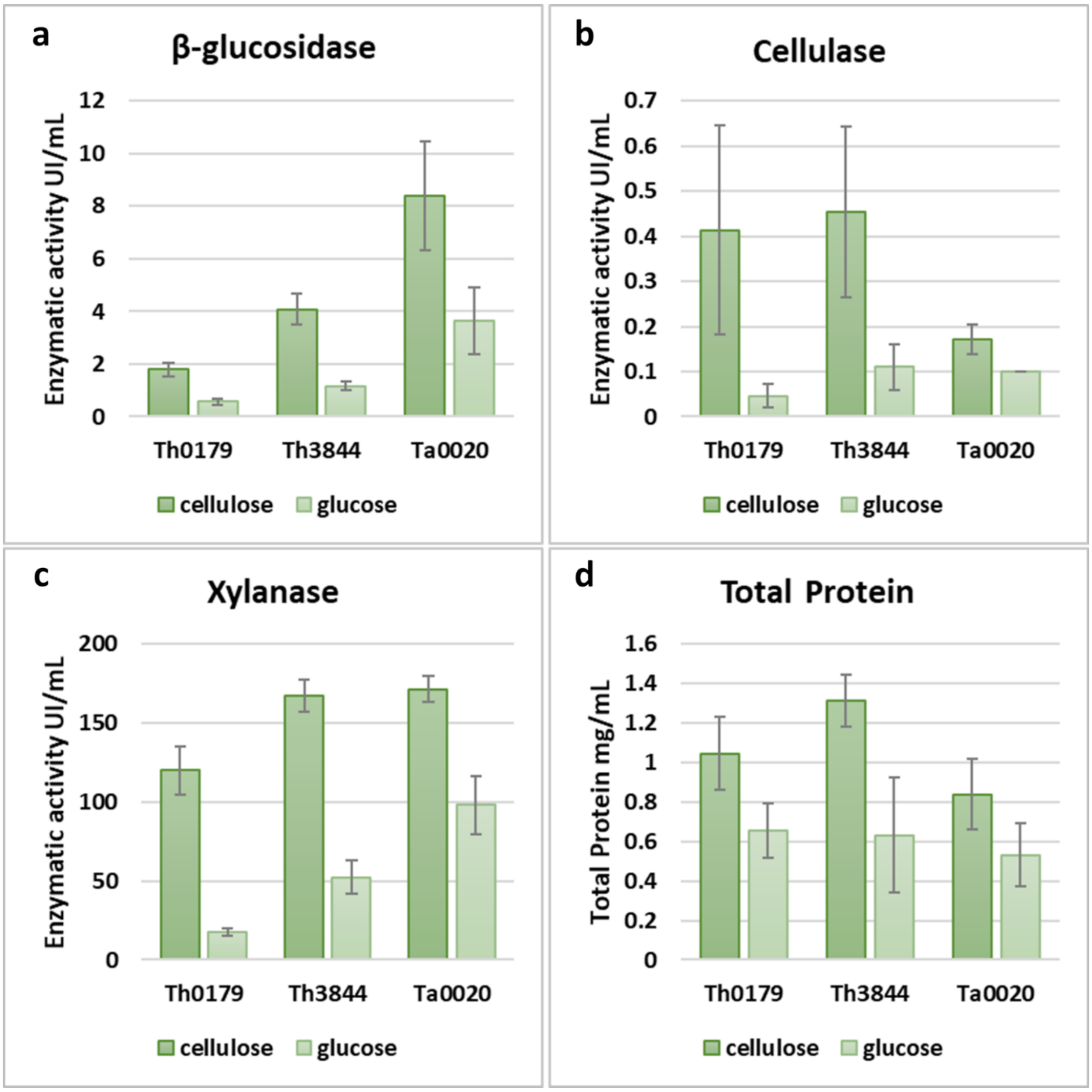


**Fig. S1** Enzymatic activity. Enzymatic activities (UI $mL^{-1}$) of β-glucosidase (a), cellulase (b), and xylanase (c) and protein contents (d) in the culture supernatants of *T. harzianum* CBMAI-0179 (Th0179), *T. harzianum* IOC-3844 (Th3844) and *T. atroviride* CBMAI-0020 (Ta0020) measured after 96 h of growth. Each bar represents the mean and standard deviation of biological triplicates.

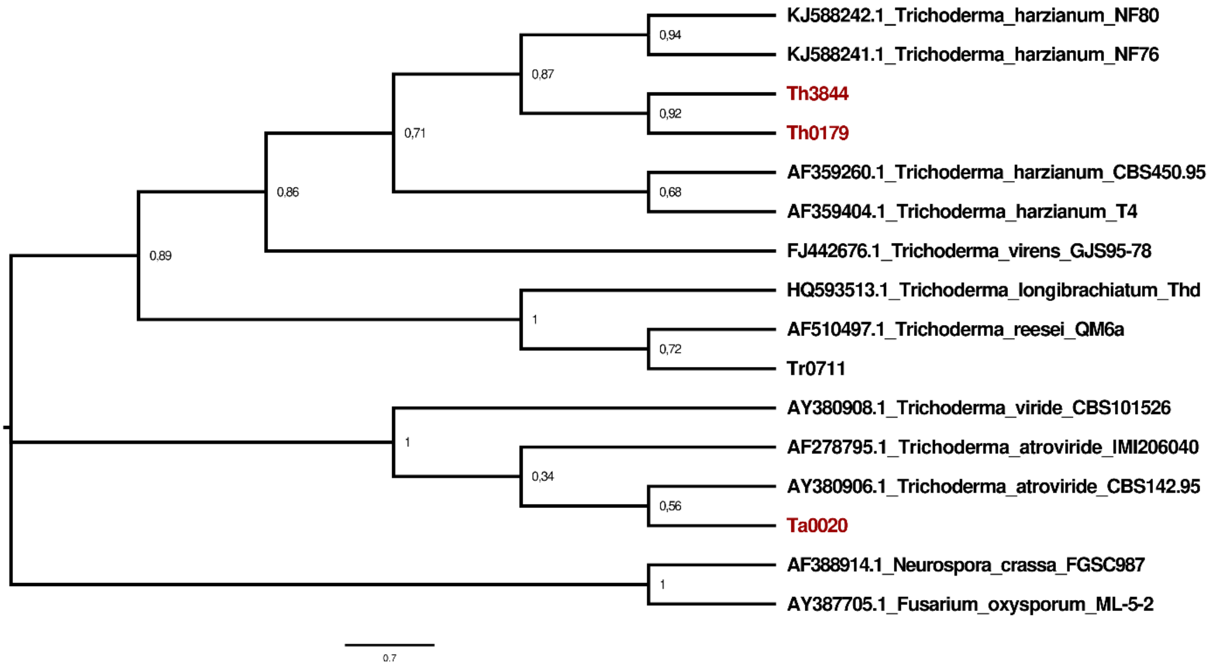


**Fig. S2** Phylogenetic tree of *Trichoderma* spp. (Rosolen et al. 2020). The sequences from the internal transcribed spacer (ITS) region were used to analyze and compare the phylogenetic relationships of *T. harzianum* CBMAI-0179 (Th0179), *T. harzianum* IOC-3844 (Th3844) and *T. atroviride* CBMAI-0020 (Ta0020). The other ITS sequences were derived from the NCBI database.

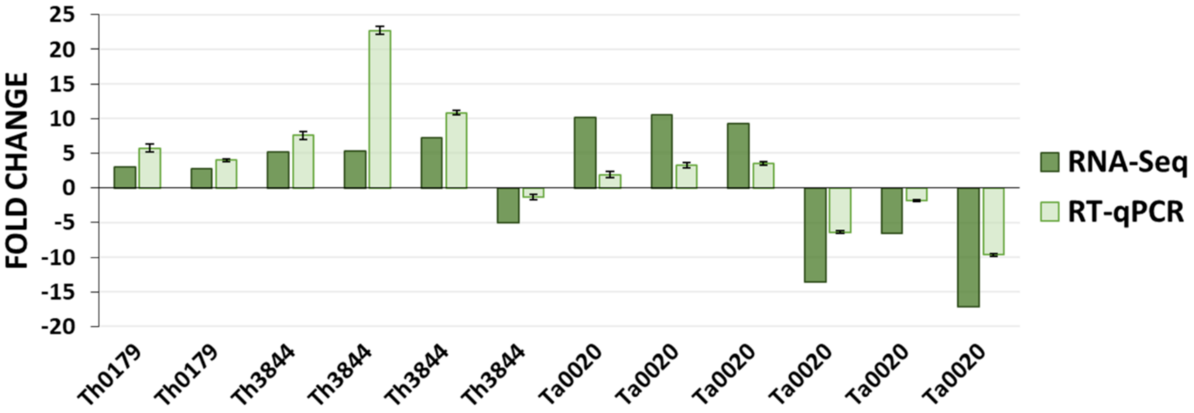


**Fig. S3** RNA-Seq analysis validation. The obtained RT-qPCR results were compared with the RNA-Seq results for transcriptome analysis validation of *T. harzianum* CBMAI-0179 (Th0179), *T. harzianum* IOC-3844 (Th3844) and *T. atroviride* CBMAI-0020 (Ta0020) using a subset of differentially expressed genes (DEGs) under cellulose or glucose growth conditions.

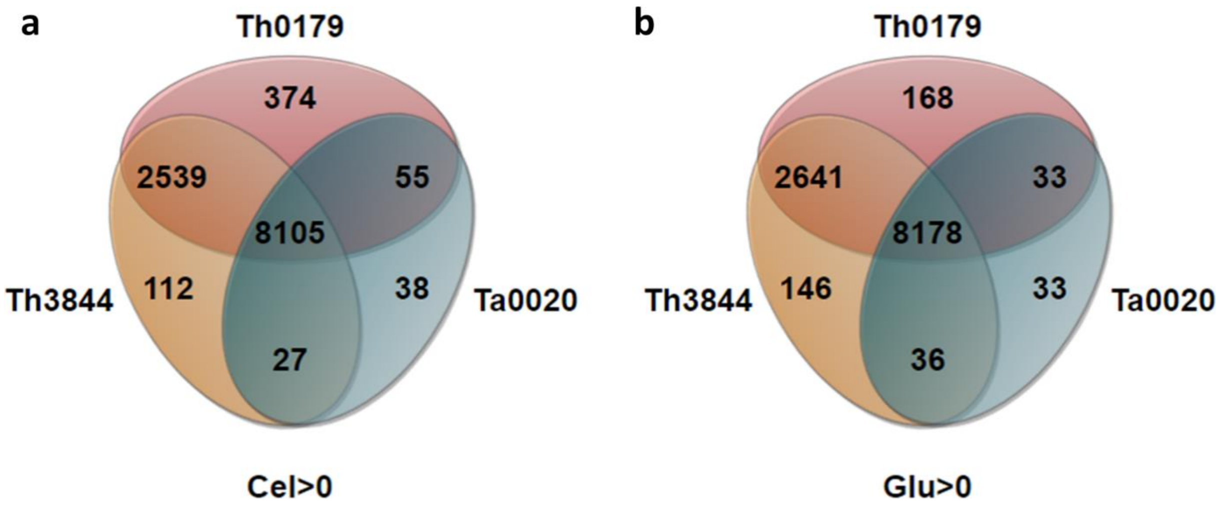


**Fig. S4** Venn diagrams. Venn diagrams of the genes identified in *T. harzianum* CBMAI-0179 (Th0179), *T. harzianum* IOC-3844 (Th3844) and *T. atroviride* CBMAI-0020 (Ta0020) with expression levels higher than zero under cellulose (Cel) (a) and glucose (Glu) (b) growth conditions using the *T. harzianum* T6776 genome as a reference.
