## Supplementary material for "The synergistic actions of hydrolytic genes reveal the mechanism of *Trichoderma harzianum* for cellulose degradation": Table S1

**Table S1** Primer sequences and amplicons of the endogenous genes evaluated in this study and differentially expressed genes (DEGs) obtained from RNA-Seq data for transcriptome analysis validation by RT-qPCR

| Gene ID | Strain | Primer Sequence (5' – 3') | Amplicon Length (bp) | Fold Change | Efficiency (%) | $R^2$ | Gene Description | E-value |
|---|---|---|---|---|---|---|---|---|
| *DEGs under cellulose or glucose conditions* | | | | | | | | |
| THAR02_01971 | Th0179 | F-CCCGTACAACAGCTCGGTAT<br>R-CCTGTGGATGTGTACGAACG | 197 | 5.74 | 98.3 | 0.99 | hexokinase | 0 |
| THAR02_02241 | Th0179 | F-CACAGCGACGAGGATGACTA<br>R-ATTAGGGCGACTTGAGAGCA | 144 | 3.99 | 104.6 | 0.99 | hypothetical protein THAR02_02241 | 0 |
| THAR02_02134 | Th3844 | F-CGTCTGCCACAAGAATGCTA<br>R-CAAGCGTCGTCTTATCCACA | 167 | 7.56 | 95.7 | 0.98 | glycosyl hydrolase family 61-2 | 0 |
| THAR02_03271 | Th3844 | F-GCCACTGATCAGGGTTCACT<br>R-CATGTTGCTGGGCATAATTG | 169 | 22.69 | 105.1 | 0.99 | glycosyl hydrolase family 10 | 0 |
| THAR02_10113 | Th3844 | F-AACCAAGGCAACAAGTGGAC<br>R-GGGTAGTTCTGCATGCCATT | 122 | 10.85 | 105.5 | 0.99 | glycosyl hydrolase family 61 | 0 |
| THAR02_08208 | Th3844 | F-GGCAACACGGAATCTCATTT<br>R-CCTTGTCGCTCGTCCTAAAG | 200 | -1.29 | 100.7 | 0.98 | hypothetical protein THAR02_08208 | 2.3E-15 |
| TRIATDRAFT_301285 | Ta0020 | F-CCATCTGCAACATGTCCATC<br>R-CAACAGCTCTGTCCCACTCA | 132 | 1.89 | 99.5 | 0.99 | hypothetical protein TRIATDRAFT_301285 | 0 |
| TRIATDRAFT_94197 | Ta0020 | F-TAGCTTGGGAGCGACAAGAT<br>R-CACTGACGCCCAAGGATATT | 195 | 3.3 | 94.1 | 0.98 | hypothetical protein TRIATDRAFT_94197 | 0 |
| TRIATDRAFT_83884 | Ta0020 | F-TGCAAAAGGCAAAGACACTG<br>R-TGACCTGTGAAGCAAAGACG | 141 | 3.54 | 99.5 | 0.99 | hypothetical protein TRIATDRAFT_83884 | 0 |
| TRIATDRAFT_297813 | Ta0020 | F-CTCCGTCGATAACACCGTCT<br>R-CCCTCAAGAGCTTGCCATAG | 152 | -6.37 | 99.9 | 0.99 | hypothetical protein TRIATDRAFT_297813 | 0 |
| TRIATDRAFT_262275 | Ta0020 | F-CGAACACTCGCATACGAAGA<br>R-TAGCCCGTTGCATAATACCC | 126 | -1.81 | 96.6 | 0.99 | hypothetical protein TRIATDRAFT_262275 | 0 |
| TRIATDRAFT_189857 | Ta0020 | F-GGTTTGGGTCGTCTTCAAAA<br>R-CCTTTCTCTTCAGCCATTCG | 170 | -9.67 | 99.2 | 0.99 | hypothetical protein TRIATDRAFT_189857 | 0 |

| Gene ID | Strain | Primer Sequence (5' – 3') | Amplicon Length (bp) | Tm (°C) | Efficiency (%) | $R^2$ | Gene Description | E-value |
|---|---|---|---|---|---|---|---|---|
| *Endogenous genes* | | | | | | | | |
| THAR02_00249 | Th0179 | F-CAGCTCAGGATCTCACCACA R-CATCGTAGAGCAGACGGTCA | 167 | 60 | 108.1 | 0.98 | NAD+ synthase (glutamine-hydrolysing) | 8.3E-01 |
| THAR02_06363 | Th3844 | F-TGCCTGAACTGGCCTCTACT R-AAATCCCATCCCTGAAATCC | 183 | 60 | 103.1 | 0.96 | hypothetical protein THAR02_06363 | 9.1E-01 |
| TRIATDRAFT_161557 | Ta0020 | F-ATGAGCCTTACGAGCAGCAT R-GTGGGCTCGAGAAGACACTC | 148 | 60 | 105.6 | 0.99 | hypothetical protein TRIATDRAFT_161557,partial | 7.3E-01 |
