## Supplementary material for "The synergistic actions of hydrolytic genes reveal the mechanism of *Trichoderma harzianum* for cellulose degradation": Table S4

**Table S4** Classification of the CAZyme genes under cellulose growth conditions for *T. harzianum* CBMAI-0179, *T. harzianum* IOC-3844 and *T. atroviride* CBMAI-0020

| Gene ID | Protein Product | Fold Change | E-value | CAZy Classification | Enzyme Activity | EC Number | Cellulose TPM | Glucose TPM |
|---|---|---|---|---|---|---|---|---|
| *Classification of the CAZyme genes under cellulose growth conditions for* T. harzianum *CBMAI-0179* | | | | | | | | |
| **THAR02_00068** | KKP07860.1 | 1.87 | 1.00E-143 | GH18 CBM1 | chitinase | 3.2.1.14 | 42.13 | 22.14 |
| **THAR02_00890** | KKP07011.1 | 2.10 | 4.00E-15 | GH3 | beta-glucosidase | 3.2.1.21 | 68.64 | 32.01 |
| **THAR02_01434** | KKP06476.1 | 1.75 | 0 | GH16 | endo-1,3(4)-beta-glucanase | 3.2.1.6 | 341.65 | 191.83 |
| **THAR02_01911** | KKP05955.1 | 1.64 | 2.00E-53 | GT4 | 1-acyl-sn-glycerol-3-phosphate acyltransferase | - | 94.76 | 56.86 |
| **THAR02_02132** | KKP05758.1 | 2.89 | 0 | GH3 | beta-glucosidase | 3.2.1.21 | 16.39 | 5.56 |
| **THAR02_02133** | KKP05759.1 | 2.27 | 3.00E-160 | CBM1 | cellulase | 3.2.1.4 | 47.86 | 20.67 |
| **THAR02_02134** | KKP05760.1 | 2.60 | 0 | AA9 CBM1 | cellulase | 3.2.1.4 | 138.62 | 52.35 |
| **THAR02_02251** | KKP05610.1 | 2.21 | 0 | GH1 | beta-glucosidase | 3.2.1.21 | 88.05 | 39.02 |
| **THAR02_02560** | KKP05371.1 | 1.97 | 2.00E-29 | GH18 | uncharacterized protein | - | 151.83 | 75.55 |
| **THAR02_02979** | KKP04958.1 | 2.05 | 3.00E-114 | GH45 CBM1 | cellulase | 3.2.1.4 | 52.97 | 25.32 |
| **THAR02_03008** | KKP04907.1 | 2.02 | 7.00E-30 | GH55 | glucan 1,3-beta-glucosidase | 3.2.1.58 | 28.88 | 14.01 |
| **THAR02_03217** | KKP04674.1 | 1.52 | 8.00E-14 | GH17 | hypothetical protein THAR02_03217 | - | 71.68 | 46.17 |
| **THAR02_03271** | KKP04658.1 | 2.25 | 5.00E-137 | GH10 | endo-1,4-beta-xylanase | 3.2.1.8 | 47.58 | 20.75 |
| **THAR02_03302** | KKP04612.1 | 1.85 | 4.00E-95 | GH16 | glucan endo-1,3-beta-D-glucosidase | 3.2.1.39 | 212.11 | 112.64 |
| **THAR02_04021** | KKP03872.1 | 2.11 | 2.00E-84 | GH72 | 1,3-beta-glucanosyltransferase | 2.4.1.- | 178.64 | 82.96 |
| **THAR02_04405** | KKP03485.1 | 2.09 | 0 | GH5 CBM1 | cellulase | 3.2.1.4 | 39.92 | 18.77 |
| **THAR02_04414** | KKP03494.1 | 1.52 | 0 | GH6 CBM1 | cellulose 1,4-beta-cellobiosidase (nonreducing end) | 3.2.1.91 | 299.66 | 192.97 |

| | | | | | | | | |
|---|---|---|---|---|---|---|---|---|
| **THAR02_05432** | KKP02477.1 | 2.64 | 0 | GH1 | beta-glucosidase | 3.2.1.21 | 125.51 | 46.73 |
| **THAR02_05677** | KKP02215.1 | 40.48 | 2.00E-127 | AA7 | UDP-N-acetylmuramate dehydrogenase | - | 20.74 | 0.50 |
| **THAR02_07531** | KKP00372.1 | 2.05 | 0 | GH20 | beta-N-acetylhexosaminidase | 3.2.1.52 | 1925.74 | 920.50 |
| **THAR02_07716** | KKP00192.1 | 1.98 | 4.00E-92 | GH64 CBM6 | glucanase B | - | 271.18 | 134.28 |
| **THAR02_07777** | KKP00125.1 | 1.79 | 0 | GH18 CBM1 | chitinase | 3.2.1.14 | 108.01 | 59.24 |
| **THAR02_07958** | KKO99924.1 | 2.60 | 3.00E-162 | GH27 | alpha-galactosidase | 3.2.1.22 | 22.70 | 8.58 |
| **THAR02_08897** | KKO99004.1 | 1.73 | 0 | GH7 CBM1 | cellulose 1,4-beta-cellobiosidase (reducing end) | 3.2.1.176 | 369.92 | 209.24 |
| **THAR02_10108** | KKO97789.1 | 1.51 | 4.00E-29 | CE9 | glucosamine-6-phosphate isomerase | - | 3705.87 | 2411.20 |
| **THAR02_10110** | KKO97791.1 | 1.64 | 0 | CE9 | N-acetylglucosamine-6-phosphate deacetylase | 3.5.1.25 | 6362.40 | 3809.76 |
| **THAR02_10273** | KKO97625.1 | 1.62 | 0 | GH17 | glucan endo-1,3-beta-D-glucosidase | 3.2.1.39 | 95.61 | 57.93 |

| Gene ID | Protein Product | Fold Change | E-value | CAZy Classification | Enzyme Activity | EC Number | Cellulose TPM | Glucose TPM |
|---|---|---|---|---|---|---|---|---|
| *Classification of the CAZyme genes under cellulose growth conditions for* T. harzianum *IOC-3844* | | | | | | | | |
| **THAR02_00574** | KKP07377.1 | 2.48 | 2.00E-135 | GH93 | non-reducing end alpha-L-arabinofuranosidase | 3.2.1.55 | 21.50 | 8.79 |
| **THAR02_00630** | KKP07306.1 | 1.62 | 2.00E-25 | GH23 | hypothetical protein THAR02_00630 | - | 114.40 | 71.54 |
| **THAR02_00656** | KKP07226.1 | 2.39 | 0 | GH3 | beta-glucosidase | 3.2.1.21 | 42.13 | 17.92 |
| **THAR02_00746** | KKP07210.1 | 9.35 | 2.00E-18 | GT51 | 3-oxoacyl-[acyl-carrier-protein]reductase | 1.1.1.100 | 68.64 | 7.45 |
| **THAR02_00890** | KKP07011.1 | 2.99 | 4.00E-15 | GH3 | beta-glucosidase | 3.2.1.21 | 211.94 | 71.88 |
| **THAR02_00891** | KKP07012.1 | 2.56 | 0 | GH3 | xylan 1,4-beta-xylosidase | 3.2.1.37 | 102.19 | 40.54 |
| **THAR02_01069** | KKP06872.1 | 2.13 | 0 | GH64 | hypothetical protein THAR02_01069 | - | 231.30 | 110.52 |
| **THAR02_01195** | KKP06706.1 | 4.58 | 1.00E-36 | GH128 | hypothetical protein THAR02_01195 | - | 10.12 | 2.25 |
| **THAR02_01348** | KKP06557.1 | 1.63 | 0 | GT2 | chitin synthase | 2.4.1.16 | 147.71 | 91.93 |
| **THAR02_01449** | KKP06491.1 | 4.39 | 5.00E-155 | CE5 CBM1 | acetyl xylan esterase | 3.1.1.72 | 199.44 | 46.11 |
| **THAR02_01501** | KKP06353.1 | 5.30 | 4.00E-151 | GH12 | cellulase | 3.2.1.4 | 241.36 | 46.22 |
| **THAR02_01764** | KKP06161.1 | 2.28 | 0 | GH30_5 | galactan endo-1,6-beta-galactosidase | 3.2.1.164 | 40.21 | 17.87 |
| **THAR02_01982** | KKP05943.1 | 1.99 | 0 | GH35 | beta-galactosidase | 3.2.1.23 | 32.94 | 16.82 |
| **THAR02_02063** | KKP05866.1 | 1.51 | 6.00E-25 | GT2 | NADH dehydrogenase | 1.6.99.3 | 137.14 | 91.93 |
| **THAR02_02133** | KKP05759.1 | 6.57 | 3.00E-160 | CBM1 | cellulase | 3.2.1.4 | 167.08 | 25.83 |
| **THAR02_02134** | KKP05760.1 | 5.23 | 0 | AA9 CBM1 | cellulase | 3.2.1.4 | 580.73 | 112.80 |
| **THAR02_02147** | KKP05773.1 | 3.30 | 2.00E-128 | GH11 CBM1 | endo-1,4-beta-xylanase | 3.2.1.8 | 737.49 | 226.98 |
| **THAR02_02152** | KKP05778.1 | 5.36 | 0 | GH54 CBM42 | non-reducing end alpha-L-arabinofuranosidase | 3.2.1.55 | 23.35 | 4.42 |
| **THAR02_02251** | KKP05610.1 | 1.93 | 0 | GH1 | beta-glucosidase | 3.2.1.21 | 240.78 | 126.43 |
| **THAR02_02979** | KKP04958.1 | 3.70 | 3.00E-114 | GH45 CBM1 | cellulase | 3.2.1.4 | 176.03 | 48.33 |
| **THAR02_03008** | KKP04907.1 | 3.11 | 7.00E-30 | GH55 | glucan 1,3-beta-glucosidase | 3.2.1.58 | 27.88 | 9.10 |
| **THAR02_03127** | KKP04790.1 | 1.52 | 9.00E-133 | AA11 | hypothetical protein THAR02_03127 | - | 82.34 | 54.84 |
| **THAR02_03217** | KKP04674.1 | 2.64 | 8.00E-14 | GH17 | hypothetical protein THAR02_03217 | - | 43.42 | 16.73 |

| | | | | | | | | |
|---|---|---|---|---|---|---|---|---|
| **THAR02_03271** | KKP04658.1 | 5.30 | 5.00E-137 | GH10 | endo-1,4-beta-xylanase | 3.2.1.8 | 741.35 | 141.93 |
| **THAR02_03302** | KKP04612.1 | 1.83 | 4.00E-95 | GH16 | glucan endo-1,3-beta-D-glucosidase | 3.2.1.39 | 188.18 | 104.14 |
| **THAR02_03357** | KKP04531.1 | 3.81 | 0 | GH7 CBM1 | glycosyl hydrolase family 7-1 | 3.2.1.- | 179.79 | 47.92 |
| **THAR02_03790** | KKP04131.1 | 1.69 | 5.00E-90 | GH76 | mannan endo-1,6-alpha-mannosidase | 3.2.1.101 | 115.01 | 68.96 |
| **THAR02_03851** | KKP04059.1 | 3.19 | 0 | GH5 CBM1 | mannan endo-1,4-beta-mannosidase | 3.2.1.78 | 460.13 | 146.29 |
| **THAR02_03852** | KKP04060.1 | 3.60 | 6.00E-73 | CE8 | pectin esterase | 3.1.1.11 | 25.74 | 7.26 |
| **THAR02_04021** | KKP03872.1 | 1.95 | 2.00E-84 | GH72 | hypothetical protein THAR02_04021 | 2.4.1 | 374.07 | 194.56 |
| **THAR02_04082** | KKP03811.1 | 3.78 | 3.00E-159 | GH62 CBM1 | glycosyl hydrolase family 62 | - | 116.18 | 31.23 |
| **THAR02_04300** | KKP03616.1 | 1.55 | 0 | GH2 | beta-mannosidase | 3.2.1.25 | 79.96 | 52.33 |
| **THAR02_04344** | KKP03537.1 | 2.52 | 0 | GH71 CBM24 | glucan 1,3-alpha-glucosidase | 3.2.1.84 | 26.16 | 10.54 |
| **THAR02_04405** | KKP03485.1 | 3.85 | 0 | GH5 CBM1 | cellulase | 3.2.1.4 | 63.50 | 16.77 |
| **THAR02_04414** | KKP03494.1 | 3.92 | 0 | GH6 CBM1 | cellulose 1,4-beta-cellobiosidase(non-reducingend) | 3.2.1.91 | 986.25 | 255.29 |
| **THAR02_04912** | KKP02981.1 | 1.93 | 0 | GH18 CBM18 | chitinase | 3.2.1.14 | 141.79 | 74.75 |
| **THAR02_05432** | KKP02477.1 | 1.79 | 0 | GH1 | beta-glucosidase | 3.2.1.21 | 339.42 | 192.17 |
| **THAR02_05677** | KKP02215.1 | 3.90 | 2.00E-127 | AA7 | UDP-N-acetylmuramatedehydrogenase | 1.3.1.98 | 661.64 | 172.48 |
| **THAR02_05896** | KKP02016.1 | 9.11 | 6.00E-156 | GH11 | endo-1,4-beta-xylanase | 3.2.1.8 | 130.01 | 14.49 |
| **THAR02_06250** | KKP01661.1 | 3.60 | 0 | GH62 | non-reducing end alpha-L-arabinofuranosidase | 3.2.1.55 | 71.36 | 20.12 |
| **THAR02_06252** | KKP01663.1 | 1.97 | 0 | GH71 CBM24 | glucan 1,3-alpha-glucosidase | 3.2.1.84 | 507.72 | 261.19 |
| **THAR02_07549** | KKP00338.1 | 2.10 | 0 | GH30_3 | glucan endo-1,6-beta-glucosidase | 3.2.1.75 | 61.63 | 29.75 |
| **THAR02_07663** | KKP00250.1 | 4.99 | 4.00E-107 | CE5 CBM1 | acetyl xylan esterase | 3.1.1.72 | 28.13 | 5.72 |
| **THAR02_08235** | KKO99651.1 | 1.77 | 0 | GH18 | mannosyl-glycoproteinendo-beta-N-acetylglucosaminidase | 3.2.1.96 | 818.92 | 468.88 |
| **THAR02_08478** | KKO99423.1 | 3.46 | 0 | GH30_7 | glucosylceramidase | 3.2.1.45 | 35.53 | 10.43 |
| **THAR02_08479** | KKO99424.1 | 4.54 | 0 | CBM1 | hypothetical protein THAR02_08479 | - | 66.13 | 14.80 |

| | | | | | | | | |
|---|---|---|---|---|---|---|---|---|
| **THAR02_08630** | KKO99257.1 | 5.22 | 4.00E-138 | GH11 | endo-1,4-beta-xylanase | 3.2.1.8 | 21.54 | 4.19 |
| **THAR02_08762** | KKO99128.1 | 4.03 | 7.00E-84 | CBM13 | endo-1,4-beta-xylanase | 3.2.1.8 | 297.81 | 75.09 |
| **THAR02_08858** | KKO99033.1 | 3.52 | 4.00E-134 | GH11 | endo-1,4-beta-xylanase | 3.2.1.8 | 124.58 | 35.91 |
| **THAR02_08897** | KKO99004.1 | 3.42 | 0 | GH7<br>CBM1 | cellulose 1,4-beta-cellobiosidase(reducingend) | 3.2.1.176 | 1697.10 | 504.35 |
| **THAR02_09385** | KKO98503.1 | 1.60 | 8.00E-59 | GH16 | endo-1,3(4)-beta-glucanase | 3.2.1.6 | 199.41 | 126.26 |
| **THAR02_09460** | KKO98433.1 | 1.84 | 0 | GH71<br>CBM24 | glucan endo-1,3-alpha-glucosidase | 3.2.1.59 | 331.69 | 182.63 |
| **THAR02_09584** | KKO98320.1 | 2.37 | 1.00E-41 | CE6 | triacylglycerol lipase | 3.1.1.3 | 60.58 | 25.94 |
| **THAR02_09719** | KKO98175.1 | 3.22 | 0 | GH5<br>CBM1 | cellulase | 3.2.1.4 | 165.87 | 52.31 |
| **THAR02_10113** | KKO97781.1 | 7.18 | 2.00E-162 | AA9<br>CBM1 | cellulase | 3.2.1.4 | 262.90 | 37.17 |
| **THAR02_10453** | KKO97447.1 | 1.73 | 6.00E-173 | CE5 | cutinase | 3.1.1.74 | 225.72 | 132.57 |
| **THAR02_11082** | KKO96812.1 | 3.47 | 0 | AA1_2 | laccase | 1.10.3.2 | 17.43 | 5.11 |
| **THAR02_11277** | KKO96621.1 | 5.22 | 0 | GH30_7 | cellulosome enzyme | - | 37.21 | 7.23 |
| **THAR02_11278** | KKO96622.1 | 3.73 | 0 | CE15<br>CBM1 | DNA topoisomerase | 5.99.1.2 | 54.53 | 14.83 |

| Gene ID | Protein Product | Fold Change | E-value | CAZy Classification | Enzyme Activity | EC Number | Cellulose TPM | Glucose TPM |
|---|---|---|---|---|---|---|---|---|
| *Classification of the CAZyme genes under cellulose growth conditions for* T. atroviride *CBMAI-0020* | | | | | | | | |
| **TRIATDRAFT_48371** | 013937547.1 | 4.67 | 2.00E-57 | GH0 | glucan 1,3-beta-glucosidase | 3.2.1.58 | 8.38 | 1.78 |
| **TRIATDRAFT_159436** | 013938409.1 | 2.89 | 2.00E-110 | GT2 | triacylglycerol lipase | 3.1.1.3 | 406.75 | 139.27 |
| **TRIATDRAFT_153371** | 013938570.1 | 1.63 | 5.00E-157 | GT4 | thioredoxin-disulfide reductase | 1.8.1.9 | 68.14 | 41.34 |
| **TRIATDRAFT_301497** | 013939152.1 | 2.09 | 3.00E-41 | GH2 | phytanoyl-CoA dioxygenase | 1.14.11.18 | 155.46 | 73.59 |
| **TRIATDRAFT_286660** | 013939498.1 | 3.53 | 5.00E-164 | CE4 | homoserine dehydrogenase | 1.1.1.3 | 153.60 | 43.07 |
| **TRIATDRAFT_302343** | 013940290.1 | 1.53 | 3.00E-83 | GH20 | phosphate-transporting ATPase | 3.6.3.27 | 58.69 | 37.95 |
| **TRIATDRAFT_161159** | 013940821.1 | 1.77 | 0 | GH3 | xylan 1,4-beta-xylosidase | 3.2.1.37 | 37.97 | 21.29 |
| **TRIATDRAFT_301356** | 013941758.1 | 2.25 | 9.00E-93 | GH13_11 | Glycine hydroxymethyltransferase | 2.1.2.1 | 692.63 | 304.80 |
| **TRIATDRAFT_300337** | 013942233.1 | 1.51 | 9.00E-88 | GH13_11 | Glycine hydroxymethyltransferase | 2.1.2.1 | 130.79 | 85.91 |
| **TRIATDRAFT_127833** | 013942429.1 | 3.19 | 4.00E-56 | GH36 | 2-alkenal reductase[NAD(P)+] | 1.3.1.74 | 40.61 | 12.60 |
| **TRIATDRAFT_79361** | 013942477.1 | 1.60 | 0 | CE9 | N-acetylglucosamine-6-phosphatedeacetylase | 3.5.1.25 | 785.40 | 487.25 |
| **TRIATDRAFT_257866** | 013942480.1 | 1.62 | 4.00E-30 | CE9 | glucosamine-6-phosphatedeaminase | 3.5.99.6 | 1198.26 | 732.37 |
| **TRIATDRAFT_285140** | 013942693.1 | 1.99 | 6.00E-75 | GH76 | glycoside hydrolase family 76 protein | - | 56.39 | 28.05 |
| **TRIATDRAFT_16857** | 013942791.1 | 2.20 | 6.00E-140 | GH75 | glycoside hydrolase family 75 protein, partial | - | 169.45 | 76.39 |
| **TRIATDRAFT_37969** | 013942813.1 | 1.66 | 0 | GH16 | endo-1,3(4)-beta-glucanase | 3.2.1.6 | 90.71 | 54.09 |
| **TRIATDRAFT_88379** | 013942958.1 | 1.95 | 0 | AA2 | catalase-peroxidase | 1.11.1.21 | 1192.59 | 606.15 |
| **TRIATDRAFT_300064** | 013943599.1 | 1.57 | 3.00E-45 | GH76 | alcohol dehydrogenase | 1.1.1.1 | 7998.18 | 5042.11 |
| **TRIATDRAFT_41039** | 013943791.1 | 2.58 | 0 | GH20 | beta-N-acetylhexosaminidase | 3.2.1.52 | 383.88 | 147.58 |
| **TRIATDRAFT_41194** | 013943809.1 | 2.78 | 3.00E-176 | GH64 | glucan endo-1,3-beta-D-glucosidase | 3.2.1.39 | 48.63 | 17.34 |

| | | | | | | | | |
|---|---|---|---|---|---|---|---|---|
| **TRIATDRAFT_283278** | 013943899.1 | 3.87 | 4.00E-141 | GH1 | serine-tRNA ligase | 6.1.1.11 | 405.38 | 103.85 |
| **TRIATDRAFT_217415** | 013945238.1 | 2.93 | 0 | GH18 | mannosyl-glycoprotein endo-beta-N-acetylglucosaminidase | 3.2.1.96 | 20.83 | 7.05 |
| **TRIATDRAFT_45299** | 013945550.1 | 1.61 | 4.00E-57 | GH16 | licheninase | 3.2.1.73 | 45.45 | 28.03 |
| **TRIATDRAFT_81098** | 013945935.1 | 3.00 | 0 | GH54 CBM42 | non-reducing end alpha-L-arabinofuranosidase | 3.2.1.55 | 18.07 | 5.97 |
| **TRIATDRAFT_298187** | 013946139.1 | 7.65 | 3.00E-29 | CBM5 | trypsin | 3.4.21.4 | 227.33 | 29.44 |
| **TRIATDRAFT_81867** | 013946279.1 | 1.82 | 0 | GH5_24 | cellulase | 3.2.1.4 | 48.13 | 26.27 |
| **TRIATDRAFT_281288** | 013946554.1 | 1.52 | 6.00E-96 | GT47 | adenylosuccinate synthase | 6.3.4.4 | 99.60 | 65.02 |
| **TRIATDRAFT_297494** | 013947658.1 | 2.04 | 1.00E-55 | GH78 | alpha-L-rhamnosidase | 3.2.1.40 | 232.56 | 112.97 |
| **TRIATDRAFT_254673** | 013948145.1 | 2.99 | 1.00E-18 | GH3 | gluconokinase | 2.7.1.12 | 42.08 | 13.93 |
| **TRIATDRAFT_289117** | 013948526.1 | 2.46 | 3.00E-16 | GT51 | 3-oxoacyl-[acyl-carrier-protein] reductase | 1.1.1.100 | 54.73 | 22.05 |
| **TRIATDRAFT_83315** | 013948942.1 | 2.54 | 3.00E-72 | GH93 | glycoside hydrolase family 93 protein | - | 33.64 | 13.13 |
