## Supplementary material for "The synergistic actions of hydrolytic genes reveal the mechanism of *Trichoderma harzianum* for cellulose degradation": Table S5

**Table S5** Proteins identified in both the transcriptome and exoproteome of *T. harzianum* CBMAI-0179 under cellulose, glucose or both growth conditions based on *T.harzianum* T6776 public genome

| Gene ID | Conditions | GenBank Protein Accession | Protein Name | E-value | Pident | CAZy Classification | EC Number | Cellulose TPM | Glucose TPM |
|---|---|---|---|---|---|---|---|---|---|
| **THAR02_00377** | Cellulose | G0RX84 | Predicted protein | 3.00E-15 | 44.58 | - | - | 1113.19 | 1128.71 |
| **THAR02_00656** | Cellulose | A0A0F9XRC5 | Beta-glucosidase | 0 | 100 | GH3 | 3.2.1.21 | 5.08 | 2.55 |
| **THAR02_00890** | Cellulose | A0A0F9XQT4 | Beta-glucosidase | 0 | 100 | GH3 | 3.2.1.21 | 68.64 | 32.01 |
| **THAR02_01069** | Cellulose | G9MX73 | Glycoside hydrolase family 64 protein | 1.00E-64 | 37.43 | GH64 | - | 17.67 | 25.95 |
| **THAR02_01434** | Cellulose | A0A0F9XP75 | Uncharacterized protein | 0 | 100 | GH16 | 3.2.1.6 | 341.65 | 191.83 |
| **THAR02_01982** | Cellulose | A0A0F9Y1F6 | Beta-galactosidase | 0 | 100 | GH35 | 3.2.1.23 | 10.08 | 6.85 |
| **THAR02_02133** | Cellulose | A0A0G0AME2 | Uncharacterized protein | 0 | 100 | CBM1 | - | 47.86 | 20.67 |
| **THAR02_02134** | Cellulose | A0A0F9XMI8 | Cellulase | 0 | 100 | AA9<br>CBM1 | 3.2.1.4 | 138.62 | 52.35 |
| **THAR02_02147** | Cellulose | A0A0F9Y0Y9 | Endo-1,4-beta-xylanase | 0 | 100 | GH11<br>CBM1 | 3.2.1.8 | 119.56 | 86.24 |
| **THAR02_02289** | Cellulose | A0A0F9Y0G5 | Cel74a | 0 | 100 | CBM1 | - | 22.53 | 24.27 |
| **THAR02_03210** | Cellulose | A0A0F9ZXC9 | WSC domain-containing protein | 0 | 100 | AA5_1 | - | 33.57 | 38.75 |
| **THAR02_03271** | Cellulose | A0A0F9XXA4 | Beta-xylanase | 0 | 100 | GH10 | 3.2.1.8 | 47.58 | 20.75 |
| **THAR02_03624** | Cellulose | G0RX52 | Extracellular metalloproteinase (Fungalysin) | 0 | 86.41 | CBM50 | 3.4.24.- | 1.02 | 0.31 |
| **THAR02_03851** | Cellulose | A0A0G0AGG8 | Mannan endo-1,4-β-mannosidase | 0 | 100 | GH5<br>CBM1 | 3.2.1.78 | 67.98 | 57.54 |
| **THAR02_04062** | Cellulose | A0A0F9XH17 | Uncharacterized protein | 2.00E-170 | 100 | - | - | 21.80 | 18.38 |
| **THAR02_04344** | Cellulose | G9MY63 | Glycoside hydrolase family 71 protein | 0 | 68.53 | GH71<br>CBM24 | 3.2.1.59 | 3.82 | 3.87 |
| **THAR02_04405** | Cellulose | A0A0F9XG06 | Cellulase | 0 | 100 | GH5<br>CBM1 | 3.2.1.4 | 39.92 | 18.77 |
| **THAR02_04414** | Cellulose | A0A0G0AEM7 | Cellulose 1,4-β-cellobiosidase (nonreducing end) | 0 | 100 | GH6<br>CBM1 | 3.2.1.91 | 299.66 | 192.97 |
| **THAR02_04626** | Cellulose | G9NK86 | Glycoside hydrolase family 92 protein | 0 | 65.15 | GH92 | - | 1.62 | 0.36 |
| **THAR02_04782** | Cellulose | A0A024HVI0 | Chitinase 18-5 (Fragment) | 6.00E-80 | 46.13 | GH18 | - | 21.24 | 16.93 |

| | | | | | | | | | |
|---|---|---|---|---|---|---|---|---|---|
| **THAR02_05380** | Cellulose | A0A0F9XQN9 | Uncharacterized protein | 3.00E-171 | 100 | - | - | 102.99 | 81.78 |
| **THAR02_05501** | Cellulose | G0R911 | Glycoside hydrolase family 92 | 0 | 54.4 | GH92 | - | 3.88 | 2.03 |
| **THAR02_05896** | Cellulose | A0A0H3UCP8 | Endo-1,4-beta-xylanase | 8.00E-156 | 98.64 | GH11 | 3.2.1.8 | 33.19 | 27.46 |
| **THAR02_06252** | Cellulose | A0A0F9XN06 | Murein transglycosylase | 0 | 100 | GH71 CBM24 | 3.2.1.59 | 81.33 | 131.74 |
| **THAR02_07321** | Cellulose | A0A0F9X7S7 | Uncharacterized protein | 0 | 100 | - | - | 33.34 | 42.55 |
| **THAR02_07975** | Cellulose | G0RXE3 | Predicted protein | 0 | 86.74 | - | - | 8.39 | 15.64 |
| **THAR02_08235** | Cellulose | A0A0F9ZHA7 | Chitinase 3 | 0 | 100 | GH18 | 3.2.1.96 | 62.16 | 62.00 |
| **THAR02_08478** | Cellulose | A0A0G0A296 | Uncharacterized protein | 0 | 100 | GH30_7 | 3.2.1.- | 6.37 | 2.67 |
| **THAR02_08479** | Cellulose | A0A0F9X463 | Uncharacterized protein | 0 | 100 | CBM1 | - | 28.08 | 15.80 |
| **THAR02_09247** | Cellulose | A0A0F9ZZN6 | Uncharacterized protein | 0 | 100 | CBM43 | - | 142.27 | 133.87 |
| **THAR02_09257** | Cellulose | E2PTX8 | Endochitinase 42 (Fragment) | 6.00E-36 | 32.07 | GH18 | - | 8.28 | 8.50 |
| **THAR02_09719** | Cellulose | A0A0F9WYH5 | Cellulase | 0 | 100 | GH5 CBM1 | 3.2.1.4 | 18.31 | 12.14 |
| **THAR02_00568** | Glucose | A0A0F9XRP2 | Uncharacterized protein | 2.00E-96 | 100 | - | - | 8.72 | 15.63 |
| **THAR02_00585** | Glucose | G9MXG7 | Uncharacterized protein | 1.00E-32 | 31.01 | - | - | 727.77 | 1005.69 |
| **THAR02_00832** | Glucose | B9VRJ1 | Chitinase (Chitinase 1) (Endochitinase 42) | 0 | 100 | GH18 | 3.2.1.14 | 143.38 | 128.50 |
| **THAR02_01200** | Glucose | Q8WZM5 | Trypsin-like protease | 0 | 97.67 | CBM5 | - | 39.89 | 32.27 |
| **THAR02_04684** | Glucose | A0A0F9XF89 | Uncharacterized protein | 0 | 100 | GH55 | 3.2.1.58 | 24.99 | 18.87 |
| **THAR02_05625** | Glucose | A0A0F9XAS9 | Neutral/alkaline non-lysosomal ceramidase | 0 | 100 | - | - | 44.77 | 35.76 |
| **THAR02_05687** | Glucose | A0A0F9XCA7 | Uncharacterized protein | 0 | 100 | - | - | 1275.27 | 1076.18 |
| **THAR02_07531** | Glucose | A0A0F9ZJ74 | Beta-hexosaminidase | 0 | 100 | GH20 | 3.2.1.52 | 1925.75 | 920.50 |
| **THAR02_08129** | Glucose | A0A0F9ZHI0 | Endochitinase 1 | 0 | 100 | GH18 | - | 25.08 | 47.01 |
| **THAR02_08897** | Glucose | A0A0F9X2V9 | Glucanase | 0 | 100 | GH7 CBM1 | 3.2.1.176 | 369.92 | 209.25 |
| **THAR02_09150** | Glucose | A0A0F9X224 | Uncharacterized protein | 0 | 100 | - | - | 119.40 | 158.96 |
| **THAR02_10300** | Glucose | G9MPI0 | Uncharacterized protein | 0 | 86.92 | - | - | 3.35 | 3.95 |
| **THAR02_00025** | Cellulose and Glucose | A0A0G0A6L8 | Glycosyl hydrolase | 0 | 100 | GH92 | - | 14.95 | 10.19 |
| **THAR02_00503** | Cellulose and Glucose | A0A0F9XRW0 | Glycosyl hydrolase family 18-1 | 0 | 100 | GH18 | 3.2.1.14 | 24.33 | 65.26 |
| **THAR02_00657** | Cellulose and Glucose | A0A0G0A4Q2 | Alpha-L-arabinofuranosidase B | 0 | 100 | GH54 CBM42 | 3.2.1.55 | 13.24 | 9.51 |
| **THAR02_00891** | Cellulose and Glucose | A0A0G0A408 | Xylan 1,4-beta-xylosidase | 0 | 100 | GH3 | 3.2.1.37 | 26.51 | 15.54 |

| | | | | | | | | | |
|---|---|---|---|---|---|---|---|---|---|
| **THAR02_01431** | Cellulose and Glucose | P00330 | Alcohol dehydrogenase 1 (YADH-1) | 1.00E-56 | 35.47 | - | 1.1.1.1 | 2.85 | 3.20 |
| **THAR02_01480** | Cellulose and Glucose | A0A0G0API0 | Uncharacterized protein | 0 | 100 | - | - | 102.98 | 74.93 |
| **THAR02_01852** | Cellulose and Glucose | A0A0F9XN15 | Alpha-galactosidase (Melibiase) | 0 | 100 | GH27 | 3.2.1.22 | 13.79 | 10.91 |
| **THAR02_01871** | Cellulose and Glucose | A0A0G0AN43 | Murein transglycosylase | 0 | 100 | GH17 | - | 786.03 | 792.56 |
| **THAR02_02038** | Cellulose and Glucose | S5RDL5 | Eliciting plant response protein | 3.00E-98 | 99.28 | - | - | 1700.66 | 2009.38 |
| **THAR02_03429** | Cellulose and Glucose | A0A0F9XJ35 | Uncharacterized protein | 3.00E-165 | 100 | - | - | 1006.65 | 776.84 |
| **THAR02_03951** | Cellulose and Glucose | A0A0G0AG54 | Glycosyl hydrolase family 31 | 0 | 100 | GH31 | - | 60.01 | 39.29 |
| **THAR02_06672** | Cellulose and Glucose | A0A0G0A7Y5 | Glucoamylase (1,4-alpha-D-glucan glucohydrolase) (Glucan 1,4-alpha-glucosidase) | 0 | 100 | GH15 CBM20 | 3.2.1.3 | 60.73 | 51.75 |
| **THAR02_07588** | Cellulose and Glucose | G9N192 | Uncharacterized protein | 4.00E-43 | 59.55 | - | - | 3702.69 | 3901.64 |
| **THAR02_07716** | Cellulose and Glucose | A0A0F9ZIR5 | Glucanase B | 0 | 100 | GH64 | - | 271.18 | 134.28 |
| **THAR02_07728** | Cellulose and Glucose | A0A0F9X6H8 | Glutaminase A | 0 | 100 | CBM13 | - | 39.56 | 29.40 |
| **THAR02_07812** | Cellulose and Glucose | A0A0G0A4H5 | Uncharacterized protein | 0 | 100 | - | - | 14.68 | 25.41 |
| **THAR02_08746** | Cellulose and Glucose | A0A0F9X1Q3 | Uncharacterized protein | 0 | 100 | - | - | 433.78 | 341.91 |
| **THAR02_10248** | Cellulose and Glucose | A0A0F9ZWU1 | Carboxypeptidase A | 0 | 100 | - | - | 5.51 | 5.88 |
| **THAR02_10337** | Cellulose and Glucose | A0A0F9WYR7 | alpha-1,2-Mannosidase | 0 | 100 | GH47 | 3.2.1.113 | 84.53 | 76.65 |
| **THAR02_11278** | Cellulose and Glucose | A0A0F9X731 | Cip2 (Fragment) | 0 | 100 | CBM1 | 3.1.1.- | 28.80 | 26.71 |
